# Combining high-resolution imaging, deep learning, and dynamic modelling to separate disease and senescence in wheat canopies

**DOI:** 10.1101/2023.03.01.530609

**Authors:** Jonas Anderegg, Radek Zenkl, Achim Walter, Andreas Hund, Bruce A. McDonald

## Abstract

Maintenance of sufficient healthy green leaf area after anthesis is key to ensuring an adequate assimilate supply for grain filling. Tightly regulated age-related physiological senescence and various biotic and abiotic stressors drive overall greenness decay dynamics under field conditions. Besides direct effects on green leaf area in terms of leaf damage, stressors often anticipate or accelerate physiological senescence, which may multiply their negative impact on grain filling. Here, we present an image processing methodology that enables the monitoring of chlorosis and necrosis separately for ears and shoots (stems + leaves) based on deep learning models for semantic segmentation and color properties of vegetation. A vegetation segmentation model was trained using semi-synthetic training data generated using image composition and generative adversarial neural networks, which greatly reduced the risk of annotation uncertainties and annotation effort. Application of the models to image time-series revealed temporal patterns of greenness decay as well as the relative contributions of chlorosis and necrosis. Image-based estimation of greenness decay dynamics was highly correlated with scoring-based estimations (r ≈ 0.9). Contrasting patterns were observed for plots with different levels of foliar diseases, particularly septoria tritici blotch. Our results suggest that tracking the chlorotic and necrotic fractions separately may enable (i) a separate quantification of the contribution of biotic stress and physiological senescence on overall green leaf area dynamics and (ii) investigation of the elusive interaction between biotic stress and physiological senescence. The potentially high-throughput nature of our methodology paves the way to conducting genetic studies of disease resistance and tolerance.

## Introduction

Final crop yields are determined through a multitude of processes and events occurring throughout the growing season. Suboptimal wheat yields can be related to limitations in sink strength and source capacity, where sink strength is defined by the number of grains and their capacity to absorb assimilates, while source capacity is defined by the capability of photosynthetically active plant tissues to provide assimilates that sustain concurrent grain filling. Despite ample evidence indicating prevalent sink limitation of wheat yields under a broad range of environmental conditions (reviewed by Araus et al., 2008 and Borrás et al., 2004), reports indicating source-limitation are not uncommon. For example, the stay-green phenotype that should increase the availability of assimilates during grain filling is often positively correlated with yields, particularly under end-of-season stress conditions (Anderegg et al., 2020; Christopher et al., 2016, 2008; Joshi et al., 2007; Verma et al., 2004). Similarly, yield-reducing effects of certain foliar diseases are thought to arise primarily as a consequence of increasing source limitation during grain filling through losses of photosynthetically active green leaf area resulting from the formation of chlorotic and necrotic lesions as well as induced necrosis (e.g., Robert et al., 2006, 2005). This is the case for septoria tritici blotch (STB) caused by *Zymoseptoria tritici*, a major fungal pathogen of wheat around the world. Even though STB lesions can be found over the majority of the growing season, a long latent period can allow several new leaf layers to develop at the top of the canopy during vegetative and reproductive growth stages before splash-dispersed spores originating from lower leaf layers reach the top leaf layers and cause new symptoms. Therefore, sink formation is not typically affected by STB, meaning that crop losses should not occur before heading or even anthesis (Bancal et al., 2007). Instead, they occur primarily as a consequence of source limitations during grain filling, when losses in green leaf area due to leaf damage become substantial.

To explain the apparent contradiction between evidence indicating sink- and source-limitation in the context of biological stresses such as foliar diseases, it may be necessary (i) to precisely quantify the time point in terms of particularly sensitive crop developmental stages at which the stress appears, as well as integrate its severity over time and (ii) understand if and how the presence of the disease interferes with whole plant functioning in ways reaching beyond the direct reduction of photosynthetically active leaf area. With respect to the latter point, it has often been suggested that, in addition to reducing green leaf area in proportion to the severity of the disease, foliar diseases may anticipate and/or accelerate senescence (Anderegg et al., 2020; Bancal et al., 2015; Simón et al., 2020), possibly through a modification of the balance between nitrogen supply by the source and demand by the sink during grain filling (Simón et al., 2020). However, detailed studies on artificially inoculated potted wheat plants under greenhouse conditions found no interaction between the presence of STB and temporal patterns of physiological senescence (Bancal et al., 2016; Slimane et al., 2012). These findings seem to contradict evidence from field experiments, where a significant anticipation of the generalized end-of-season decay in measures of canopy greenness is often observed and frequently reported to be closely related to STB-related yield losses (Bancal et al., 2015).

Separating the effects of diseases and physiological senescence on overall greenness decay using visual assessments is a daunting task, even at the level of individual leaves. This also holds true for currently available assessment strategies relying on destructive sampling of plant material and subsequent image analysis, which is largely based on color properties of sampled materials (Anderegg et al., 2022, 2019; Stewart et al., 2016), that may perfectly overlap between senescent and diseased necrotic tissue. Yet, whereas the final outcome (necrosis) is the same for STB and physiological senescence, the latter is a more gradual and generalized process, typically encompassing a widespread yellowing (chlorosis) of plant tissues following controlled chlorophyll degradation. Some chlorosis is frequently observed surrounding necrotic STB lesions as well, however to a very limited spatial extent (typically less than 1% of the total leaf area; Anderegg et al., 2022). We therefore hypothesize that the occurrence and the relative contributions of chlorosis and necrosis to greenness decay may enable a quantification and a separation of the effects of (biotic) stresses and physiological senescence on the maintenance of source capacity during grain filling. However, currently available assessment strategies are either too laborious and interfere excessively with the development of epidemics when carried out frequently (destructive samplings) or are imprecise and subjective (visual scorings) which stands in contradiction with the need for temporally highly resolved data that is comparable across experiments.

Recent advances in sensor and carrier platform technology as well as in processing and analysis of resulting data sets are increasingly enabling fast and objective sensor-based quantification of various crop traits under field conditions. For example, Grieder et al. (2015) used repeated close-range RGB imagery during winter to characterize the temperature response of early canopy growth in different wheat genotypes. Recently, deep convolutional neural networks (CNNs) have proven useful for various tasks related to phenotypic trait extraction from images, including object detection and counting (e.g., of wheat ears [David et al., 2020] or sorghum panicles [James et al., 2023]) and image segmentation (e.g., vegetation-soil segmentation [Serouart et al., 2022; Zenkl et al., 2022] or ear segmentation [Dandrifosse et al., 2022b]). Serouart et al. (2022) used a deep learning model to segment vegetation from soil background and a support vector classifier to partition detected vegetation into chlorophyll-active and inactive vegetation.

Unfortunately, generating pixel-level image annotations from scratch for the training of semantic segmentation models can be extremely time consuming. In addition, if the annotation task is challenging, there is a high risk of annotation uncertainties significantly lowering the performance ceiling for segmentation models (Zenkl et al., 2022). Maximizing the quality and efficiency of image annotations is therefore a key objective when generating training data. This is true particularly when diverse application scenarios (as encountered when monitoring diverse genetic material across different phenological stages under field conditions) require a broad training and evaluation data base to adequately represent most relevant scenarios.

A key advantage of sensor-based phenotyping may lie in the possibility to accurately track even small changes in crop canopy characteristics over time at the plot level. For example, even though diseased and naturally senescent canopies appear similar at coarse optical resolution in terms of their reflectance properties at specific points in time, tracking changes in color properties over time can be informative of the causal processes underlying the loss of canopy greenness, facilitating a separate assessment of senescence-related and disease-related effects (Anderegg et al., 2019). Unfortunately, such reflectance-based approaches are limited by the fact that canopy architectural and morphological traits such as leaf angles or leaf glaucousness differ markedly across wheat genotypes while strongly affecting mixed reflectance signals. Additionally, the contribution of different components of a measured scene such as soil background or different organs (leaves, stems, and ears) change dynamically over time in a genotype-dependent manner, introducing significant bias even when only relative signal changes over time are analyzed (Anderegg et al., 2020). High-resolution image data with a pixel-resolution in the sub-millimeter range facilitates the extraction of organ-level signals as well as an elimination of background signals (e.g., Dandrifosse et al., 2022a).

The main objective of this work was to develop potentially high-throughput methods facilitating a dynamic quantification of the fraction of healthy, senescing/chlorotic, and senescent/necrotic vegetation at organ-scale from image time series. This in turn will enable (i) monitoring the effect of stresses on these fractions and their dynamics during critical stages of crop development, (ii) disentangling the effects of biotic stresses and physiological senescence on overall green leaf area dynamics after anthesis and (iii) an investigation of the elusive interaction between biotic stressors and physiological senescence under field conditions. We hypothesize that such knowledge could enable the identification of STB-tolerant genotypes, because delayed physiological senescence has been identified as a promising compensation mechanism under disease pressure (Bancal et al., 2015). Here, we present a proof of concept by applying the proposed methods to time-series of high-resolution RGB images taken in a dedicated experiment with thorough ground truthing of disease intensity and senescence dynamics through established methodologies.

## Materials and Methods

### Plant materials and experimental design

A set of sixteen registered bread wheat cultivars was grown at the ETH Research Station for Plant Sciences Lindau-Eschikon, Switzerland (47.449N, 8.682E, 520 m a.s.l.; soil type: eutric cambisol) in the wheat growing season of 2021-2022. Plots were sown on October 18, 2021, with a drill sowing machine at a blade distance of 0.125 m resulting in 400 plants m^-2^. Cultivars with similar phenology and final height but with strongly contrasting canopy architectural and morphological traits were selected for this experiment, based on data from Anderegg et al. (2021). Specifically, the set comprised an equal number of cultivars with erect and planophile flag leaves and with high and low levels of flag leaf glaucousness (Supplementary Table S1). Three cultivars were selected for each factor combination. The resulting set of twelve cultivars was complemented with a highly STB-resistant and a highly STB-susceptible cultivar, with a cultivar harboring the *Lr34* disease resistance gene that causes extensive leaf-tip necrosis, and with an awned cultivar. This selection resulted in a large variability in the physical appearance of wheat stands during grain filling (Supplementary Figure S1). Each cultivar was grown in nine plots sized 1 m × 1.7 m and one of the following three treatments was allocated to each plot with the aim of maximizing variability in STB disease severity: (i) an early fungicide application at jointing followed by artificial inoculations with a *Z. tritici* spore suspension at booting and heading (FI); (ii) a fungicide application at jointing without artificial inoculation (F0I), and (iii) neither fungicide application nor artificial inoculation (0F0I). The developmental stages were reached on 22 April 2022 for jointing (GS 31, according to BBCH scale of Lancashire et al., 1991), 16 May 2022 for booting (GS 45), and 24 May 2022 for heading (GS55), respectively. The aim of the ‘FI’ treatment was to generate symptoms of STB in otherwise healthy plots, whereas the ‘0F0I’ treatment was expected to result in plots with natural co-infections of multiple foliar diseases. The ‘F0I’ treatment represents standard agricultural practice under low disease pressure and was expected to result in healthy canopies. The fungicide used was ‘Input’ (Bayer; a mixture of spiroxamine at 300 g/L and prothioconazole at 150 g/L), with a dose of 1.25 L/ha. Preparation of the spore suspensions and field inoculations were done similarly as described in detail earlier (Anderegg et al., 2019). Briefly, 200 ml of a spore suspension with a total spore concentration of 10^6^ spores ml^-1^ was applied to each plot using a backpack sprayer. The spore suspension was supplemented with 0.1% of TWEEN 20 surfactant. Inoculum was sprayed in the evening into the wet canopy of each plot. Spore suspensions for the first and the second inoculation contained a mixture of six and nine *Z. tritici* strains, respectively, selected according to their mean virulence and reproductive potential on a large number of wheat genotypes to maximize expected diversity in symptom phenotypes. Fungal strain selection was based on data from Dutta et al. (2021). The two-factorial experimental design was generated using the functions *findblks()* and *facDiGGer()* of the R-package ‘DiGGer’ (Coombes, 2009).

**Table 1.** Searched hyperparameter space for the vegetation and ear segmentation models and determined optimal values.

| Category | Tuning parameter | Tested values | Optimal value |  |
| --- | --- | --- | --- | --- |
|  |  |  | Vegetation model | Ear model |
| **Network architecture** | Resnet encoder depth | {resnet18<sup>1</sup>, r'34, r'50, r'101} | resnet34 | resnet50 |
|  | Segmentation framework | {fpn<sup>2</sup>, unet++<sup>3</sup>, deeplabv3+<sup>4</sup>} | unet++ | deeplabv3+ |
| **Data augmentation** | Image size | {100k\|k∈{4, ..., 12}} | k=7 | k=6 |
|  | Blur kernel size (Gaussian) | {1, ..., 12} | 3 | 7 |
|  | p(color jitter) | {0.1k\|k∈{0, ..., 10}} | k=0 | k=0 |
| **Network training** | Strategy | {train<sup>5</sup>, freeze<sup>6</sup>, no_freeze<sup>7</sup>} | no_freeze | no_freeze |
|  | Batch size | {2, ..., max} | 15 (i.e., max) | 31 (i.e., max) |
|  | Learning rate | {10<sup>-5</sup>, ..., 10<sup>-1</sup>} | 0.09 | 0.066 |
|  | Momentum | {0.5, ..., 0.99} | 0.88 | 0.90 |

<sup>1</sup>He et al. (2016)
<sup>2</sup>Lin et al. (2017)
<sup>3</sup>Zhou et al. (2018)
<sup>4</sup>Chen et al. (2018)
<sup>5</sup>Encoder weights initialized randomly, optimized on the dataset
<sup>6</sup>Encoder weights initialized to pre-trained on ImageNet (Deng et al., 2009), not optimized
<sup>7</sup>Encoder weights initialized to pre-trained on ImageNet, optimized on the dataset

### Reference data collection and processing

Heading date and flag leaf and canopy greenness were assessed for all experimental plots by means of visual scorings at two-day intervals. Given that *Z. tritici* primarily infects leaves and anticipating the feasibility of extracting vegetation color properties at the organ-level, canopy greenness scorings were made with a focus on the total leaf area but neglecting stems and ears, in contrast to previous work (Anderegg et al., 2020). All visual scorings were performed by the same operator and recorded using the Field Book app (Rife and Poland, 2014).

The amount of STB in each plot was assessed on three dates (16 June 2022 i.e. 31/23 days post inoculation [dpi]; 23 June 2022 i.e. 38/30 dpi; and 29 June 2022 i.e. 44/36 dpi, referred to in the following as t1, t2 and t3, respectively) following a protocol described earlier (Anderegg et al., 2019). Briefly, visual assessments of disease incidence on flag leaves of 30 culms per plot were multiplied with a measurement of conditional severity in the form of the percentage leaf area covered by lesions (PLACL) obtained for eight detached and scanned infected flag leaves per plot, estimated using the method of Stewart et al. (2016).

### Image acquisition

A full-frame mirrorless digital camera (EOS R5, Canon Inc., Tokyo, Japan; 45 megapixel, 36 x 24 mm sensor) was mounted on a custom-made portable aluminum frame as in Grieder et al. (2015) to capture images from a nadir perspective with a fixed distance to the soil of 2.25 m (Supplementary Figure S2). Focal length of the camera zoom lens varied between 48 and 52 mm between measurement dates. Focal distance was kept constant at 1.8 m, and lens aperture was also kept constant at f/16, providing a depth of field of 1.3 m. This setup resulted in a ground sampling distance of ∼0.02 cm / pixel at ground level and ∼0.012 cm / pixel at the top of the canopy, while still providing sufficient depth of field for all objects of interest (i.e., lowest leaves to ear tips) to be in focus. The resulting field of view at ground level was approximately 1.0 x 0.66 m. Images were captured under stable light conditions, either under constant direct sunlight or under diffuse light conditions on 17 dates between heading and physiological maturity. An exposure compensation of −0.33 exposure values was used irrespective of the light conditions. This slightly reduced exposure represented a meaningful compromise between over-exposure of top-of-canopy flag leaves and ears while avoiding clipping of shadows in the lower leaf layers deeper in the canopy. Sufficient exposure of shaded lower leaf layers is an important consideration here because STB moves from lower leaves to upper leaves by splash-dispersed conidia, thus affecting lower leaf layers first. Data was recorded in raw format (.CR3) with 16 bits per color channel. Measurements were regularly completed for all 144 plots of the experiment within 1.0 – 1.5 hours and were carried out at different times of the day. For analysis, all images were converted to 8-bit portable graphics format (png).

### Training and validation datasets

Training and validation data sets were generated (i) for segmentation of plant foreground from soil background and (ii) for segmentation of wheat ears from the rest of the image. The objective of this work was to enable an accurate segmentation of images taken with our or comparable imaging set-ups. We specifically selected genotypes and imaging time points to maximize variability in terms of physical appearance of crop stands. We anticipate that both the training data generation protocol as well as the data sets themselves will be useful for ongoing initiatives aimed at assembling diverse data sets that enable the training of robust segmentation models (similar as in e.g., David et al., 2020). Below, we provide a description of the training and validation data generation process.

#### Vegetation Segmentation

We found that precisely annotating senescent or senescing vegetation in fully developed wheat canopies was extremely time consuming and challenging, especially under direct sunlight, with often low agreement between annotators. To circumvent problems with human annotation and cover the different appearances of a crop canopy during grain filling, we tested an alternative approach to human annotations. The approach relied exclusively on semi-synthetic training data generated requiring minimum human intervention (Figure 1). The two-step approach consists of (i) creating composite images of soil backgrounds and plant foregrounds that are sampled from separate original images and (ii) a subsequent domain-transfer of composite images using generative adversarial neural networks (GANs). This approach essentially reduced the need for human intervention to reviewing of automatically generated pre-segmentations.

**Figure 1.**
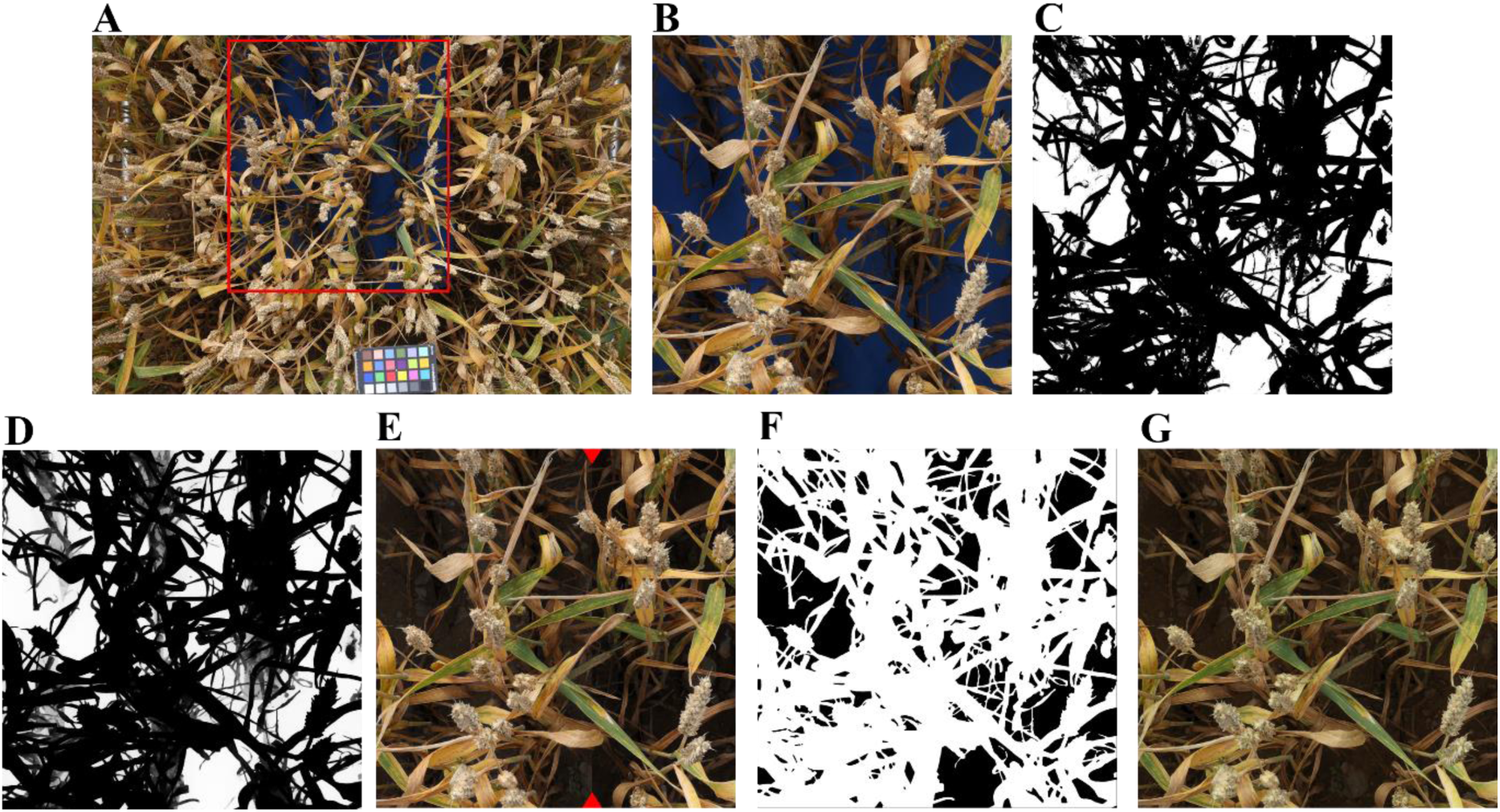
Workflow for training data generation. **(A)** Original image of a plot with the soil background covered by blue foam rubber. The red rectangle represents the identified region of interest. **(B)** A 2400 x 2400 pixels patch sampled from the region of interest in the image displayed in Figure A. In this specific case, the patch was sampled starting from the upper-right corner of the identified region of interest. **(C)** Results of the initial color-based pixel-wise segmentation. Grey-scale values represent class probabilities output by the classification model; intermediate class probabilities indicating low confidence in the classification can be found primarily in underexposed parts of the image. **(D)** Post-processed grey-scale image. **(E)** Composite images created after manual reviewing. Red arrows indicate the border between two different soil images used as background. This image was created for illustration purposes only. **(F)** The corresponding manually reviewed vegetation mask. **(G)** The domain-transferred image, used to train the vegetation segmentation model with the mask in Figure G as the target. The soil background is equivalent to the one shown in the left part of Figure E.

To generate composite images, soil in selected wheat plots was covered with readily available blue foam rubber (Rayher Hobby, Laupheim, Germany). This material could be easily placed between wheat rows, withstands weather, and is an excellent diffuse light reflector (Figure 1). Wheat plots manipulated in this way were imaged with the set-up described above under varying light conditions and at different stages of crop development. Wheat plots sown to 33 different cultivars and being part of the described experiment as well as of a neighboring wheat experiment on the same site were used. Areas in resulting images where soil was covered were then automatically cropped using simple thresholds in the HSV color space (95 ≤ H ≤ 115, 150 ≤ S ≤ 200, 22 ≤ V ≤ 255) and a single patch of 2400 x 2400 pixels was sampled from this area. These patches were then segmented pixel-wise into plant foreground and artificial background using a random forest classifier with different color spaces (RGB, L*a*b*, L*u*v*, HSV, HSI, YCbCr and YUV) as input features for each pixel. The resulting class probability maps were represented as 8-bit gray-scale images and post-processed using the Fast Bilateral Solver, an edge-aware smoothing algorithm (Barron and Poole, 2016)^1^. The post-processed probability maps were then converted into binary images by thresholding at a value of 165, which was optimized through visual inspection on a subset of images. Finally, the contours of the resulting foreground plant objects were exported as a list of polygons and imported into the Computer Vision Annotation Tool^2^ (cvat) for reviewing. Where needed, exposure was increased to ease reviewing of underexposed parts of images. A total of 206 patches of 2400 x 2400 pixels size were processed in this manner.

Each of the resulting plant foregrounds was then combined with 10 different images of bare soil which were randomly selected from a larger data base of soil images taken with the same imaging setup. Specifically, the artificial background was replaced by the corresponding areas of the selected soil image, with the light intensity pattern (gray scale values) observed on the artificial background transferred to the soil image. To compensate for the different reflectance properties of the artificial background material and natural soil, the contrast was increased by applying the simple linear transformation

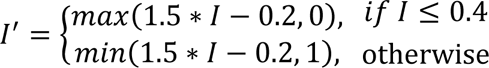

where I’ is the modified and I the original intensity value.

Although the above-described process resulted in realistic looking images in many cases, some readily recognizable differences to real images often remained, including differences in saturation between foreground and background of the composite images as well as segmentation artifacts along foreground object borders (often a faint blueish glow from mixed pixels). To resolve this issue, we performed a domain transfer using CycleGANs (Zhu et al., 2017)^3^. The overall aim of this domain transfer is to render more realistic images from the created composite images with no loss of semantic information. This is highly desirable in our case, as we wanted to avoid an additional reviewing of annotations after this step. A separate CycleGAN was trained for images captured under diffuse and under direct lighting (categorized based on date of capture). Training and validation images for both scenarios were selected to contain both healthy, green, and senescent or diseased leaf material. The models were trained using patches of 360 x 360 pixels sampled from composites and real images in original resolution. Models were trained for 200 epochs. The resulting models were used to transfer the full-sized composites (2400 x 2400 pixels), which were then used for the training of segmentation models. No real images annotated from scratch were used for training of the segmentation model.

For validation, a set of patches measuring 1200 x 1200 pixels were sampled from 74 real images and manually annotated from scratch. Images were selected to be approximately balanced with respect to date of capture, genotype, experimental treatment, STB incidence, lighting, and phenology. To reduce the annotation effort, twelve of the 16 genotypes in the experiment were sampled randomly, and some measurement dates excluded, which also reduced redundancy in the data set (e.g., from multiple measurements of the same genotype during its stay-green period).

#### Ear segmentation

Compared to the annotation of all vegetation, the annotation of spikes with well-defined borders is fast and straightforward, resulting in much lower risk of annotation errors. We therefore chose a standard approach to annotate data manually. All spikes were annotated at pixel-level in a total of 180 images using the ‘intelligent scissors’ tool of cvat. Images originated from two experiments: (i) the experiment described above, and (ii) an experiment carried out in 2015 (Grieder et al., unpublished data). Images in the 2015 experiment were acquired with a similar measurement set-up, but with a different camera and different camera settings as well as different plant material (see Grieder et al., 2015 for details).

### Segmentation model training, evaluation, and inference

Due to the different nature of the training and validation data sets for the segmentation of vegetation and ears, a separate model was trained, and model hyper-parameters were tuned separately, for each task. Tuned hyper-parameters were the depth of the ResNet encoder (He et al., 2016) and the segmentation framework, data augmentation applied (image resolution, image blurring, and the probability of applying jittering of brightness, contrast and saturation within a pre-defined range), and details of the training process (training strategy, batch size, learning rate, and momentum). Input transformations were performed using the python library ‘kornia’ (Riba et al., 2020). Random flipping, rotation, and cropping of the image were always included (i.e., not tuned). The searched parameter space and the determined optimal values are reported in Table 1. The default TPESampler of the python library ‘optuna’ (Akiba et al., 2019) was used for value suggestion. The training process was always based on a cross-entropy loss function and the stochastic gradient decent optimizer, model performance monitoring and model selection was always based on the overall validation F1-Score.

The vegetation segmentation model was evaluated on the validation data set described above. For the ear segmentation model, the available 180 annotated images were randomly split into a training and a validation data set with an 80:20 split (i.e., 144 training images and 36 validation images). Inference on all images was performed for a central region of interest sized 4000 x 4000 pixels. This cropping removed border rows from the images while keeping the central 4-5 rows. It should be noted that canopy height was not considered when cropping the images, meaning that the field of view at canopy height as well as the viewing angle distribution may differ somewhat depending on the genotype.

### Color-based classification of vegetation

The overarching goal of this work was to develop a toolset enabling the monitoring of the relative amount of healthy, chlorotic/senescing, and necrotic/senescent vegetation in time-series of images for downstream physiological studies. Hence, the final stage of the image processing workflow consisted in a classification of vegetation pixels into one of these fractions. The difference between these is in general readily observable based on color properties in-field as well as in RGB images (Anderegg et al., 2020; Cai et al., 2016; Serouart et al., 2022), although it may be difficult to define exact thresholds separating them. Here, we took an approach very similar to the one proposed by Serouart et al. (2022). Specifically, pixels making up the vegetation fraction were classified into one of the three fractions using a multiclass random forest classifier with different color spaces (RGB, L*a*b*, L*u*v*, HSV, HSI, YCbCr and YUV) as input features. In contrast to the procedure advocated by Serouart et al. (2022) for the sampling of training and validation data, we trained the model using selected patches of vegetation with an unambiguous status. This is likely to result in a biased performance estimate for the classifier, because easy-to-classify patches are preferably sampled in the process. We reasoned that a vast majority of vegetation pixels can always be confidently attributed to one of these classes, even in underexposed parts of images, with edge cases making up a very small fraction of an image. Therefore, and given the continuity between classes which renders a classification inherently subjective through the definition of arbitrary thresholds, we argue that this does not constitute a limiting factor, while greatly simplifying the generation of training data. Training data was sampled from 96 images which were selected to represent an equal number of images per genotype (n =16), per light condition (diffuse or direct sunlight) and per phenological phase during grain filling (stay-green, senescing, and senescent). Model hyper-parameters were tuned first through a randomized search to reduce the parameter search space, and subsequently through an exhaustive grid search within the reduced space. The python library ‘scikit-learn’ (Pedregosa et al., 2011) was used for this purpose. To identify the most predictive color features for this classification, we also performed feature selection using the recursive feature elimination wrapper approach, as described in detail earlier (Anderegg et al., 2020).

### Modelling of trait dynamics

Image-based time-point specific trait values were further processed to capture the dynamics of vegetation cover and vegetation status throughout the grain filling phase. Four-parameter Gompertz models were used to fit the decrease of healthy green vegetation as well as the increase in necrotic/senescent vegetation over time

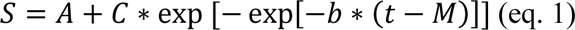

where *S* represents the trait value, *A* and (*A + C*) are the lower and upper asymptotes, respectively, *b* is the rate of change at time *M* and *M* is the time point when the rate is at its maximum (Gooding et al., 2000). Models were fitted using the R package ‘nls.multstart’ (Padfield and Matheson, 2018). Since the fraction of chlorotic plant material and vegetation cover at organ level did not show a monotonous change during the assessment period, P-splines were used for those cases. P-splines were fitted using the R package ‘scam’ (shape-constrained additive models; Pya and Wood, 2015), with the number of knots set to three quarters of the number of observations. From the fitted data, a set of dynamics parameters was then extracted (Figure 2): from the Gompertz model, all four parameters and the integral under the curve were extracted; from the P-spline fits, we extracted the maximum value, the time points when one quarter and one half of the maximum was reached in the increasing and the decaying phase (q1, q2, h1, h2) as well as the durations q2-q1 (intq) and h2-h1 (inth), and the integral under the fitted curve.

**Figure 2.**
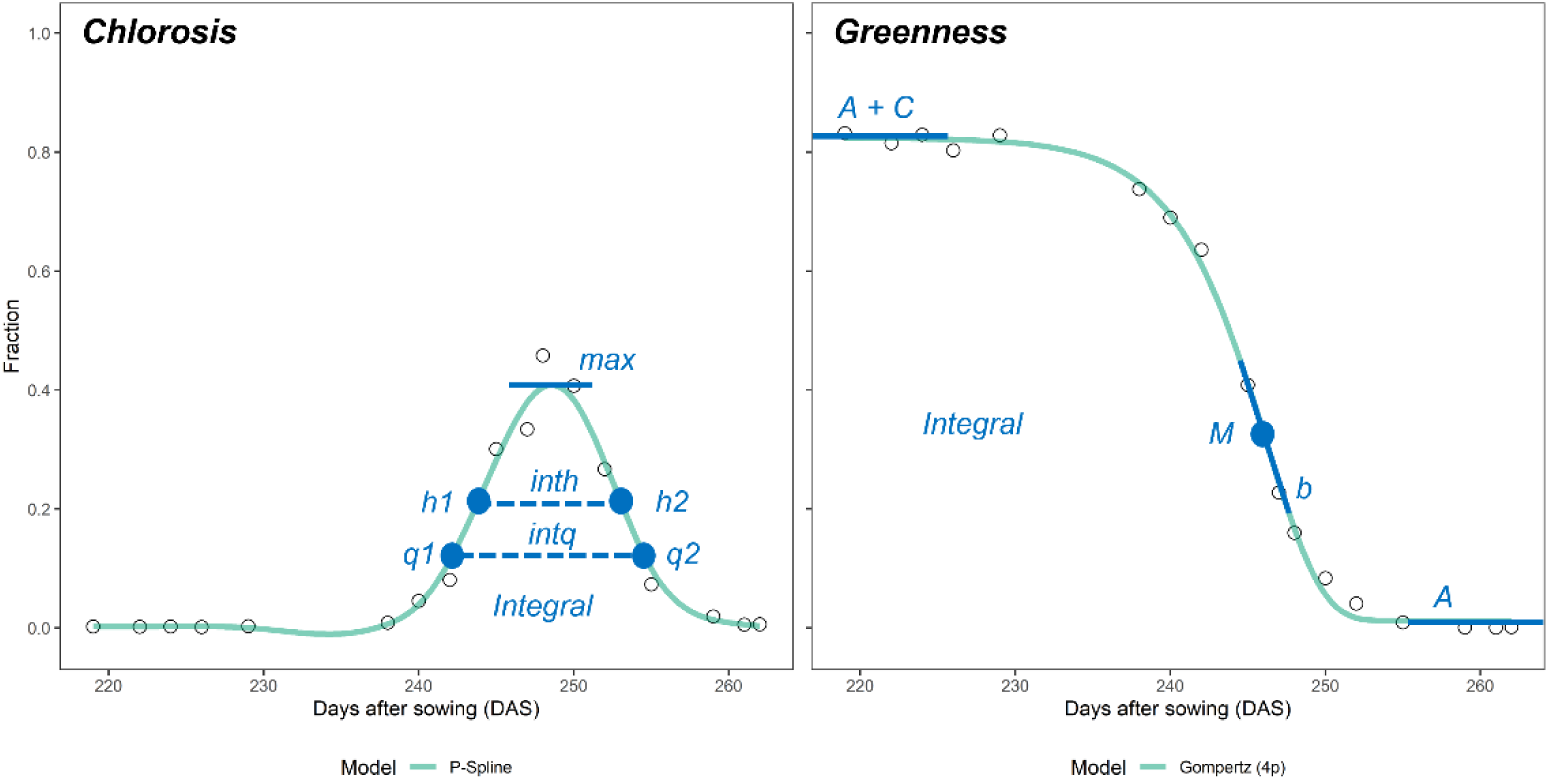
Dynamics parameters extracted for the modelled temporal trends in chlorosis and greenness extracted from a time-series of RGB images. Black circles are raw data points for one experimental plot (genotype ‘Aubusson’, treatment ‘F0I’), green lines show the model fits. Blue dots represent time-points extracted in days after sowing, blue lines indicate fractions or changes (dimensionless). For both curves, the integral was also extracted. For chlorosis, the parameter max represents the maximum of the chlorotic fraction reached; the parameters q1, q2, h1, and h2 denote the time points when one quarter and one half of the maximum was reached in the increasing and the decaying phase; the parameters intq and inth represent the durations q2 – q1 and h2 – h1, respectively. For the greenness decay, the parameters A, (A + C), b, and M are the four parameters fully describing the fitted curve, i.e., the lower and upper asymptotes, the rate of change at time M, and the time point when the rate is at its maximum, respectively.

Visual greenness scorings were linearly interpolated to daily resolution. We used linear interpolation rather than (semi-)parametric models as advocated previously (e.g., Anderegg et al., 2020; Christopher et al., 2014) because temporal patterns differed strongly across plots, likely due to the presence of treatments, which made the choice of an appropriate non-linear model difficult (Supplementary Figure S3). From the fitted data, a set of dynamics parameters were extracted: The onset, midpoint, and end of greenness decay were extracted as the time points when visual scorings fell below pre-defined thresholds (8 and 0.8, 5 and 0.5, 2 and 0.2 for visual scorings and the green fraction, respectively). The curve integrals were also extracted.

### Statistical Analysis

Spatial trends in all time-point specific and time-integrated image-based and reference traits were estimated by fitting two-dimensional P-splines to raw plot values using the R-package ‘SpATS’ v.1.0-11 (Rodríguez-Álvarez et al., 2018). Row and column were modelled as additional random effects. To obtain spatially corrected plot values and an estimation of the spatial trend, we encoded each genotype-by-treatment combination as a factor with 48 levels (i.e., one level for each of the full-factorial combinations of the 16 genotypes and 3 treatments). For the disease incidence scorings, an additional fixed effect was included in the model that specified the scorer (3 levels), with the aim of accounting for possible scorer bias. Finally, using spatially corrected plot values, treatment contrasts per trait for each genotype were extracted based on treatment means. Pearson product moment correlations between treatment contrasts in different traits were computed across the 16 genotypes included in the experiment using the function *cor.test()* of the R-package ‘stats’.

### Code and data set availability and reproducibility

All image processing and statistical analyses were implemented in python and R (R Core Team, 2018). All code and data sets pertaining to the described deep learning model optimization and training will be open-sourced on Github, an archived version will be made available *via* the ETH Zürich publications and research data repository (https://www.research-collection.ethz.ch/).

## Results

### Semi-synthetic data enabled the training of a powerful vegetation segmentation model with minimal annotation effort

Our approach for a precise and objective annotation of all vegetation in senescing and/or diseased canopies (Figure 1) enabled the generation of 206 training patches sized 2400 x 2400 pixels at the cost of approximately 80 h of annotation effort. In comparison, approximately 150 h of annotation effort were invested in the annotation from scratch of 74 validation patches sized 1200 x 1200 pixels. This represents a 20-fold decrease in time spent on annotation. Our approach also limited annotation uncertainties to parts of images where reviewing was necessary, which regarded mostly underexposed parts of images, thus ensuring high-quality annotations.

The vegetation segmentation model trained directly on the raw composite images achieved an overall validation F1-Score of 0.929, whereas the same model trained on style-transferred composite images achieved a validation F1-Score of 0.951 (Supplementary Figure S4). Thus, the CycleGAN-mediated style transfer decreased the error rate by approximately 30%. Whereas the performance of the model trained on style-transferred composites remained stable throughout the training process (validation F1-Score of approximately 0.945), the performance of models trained on raw composites tended to deteriorate as training progressed (validation F1-Score decreasing to below 0.90; Supplementary Figure S4), suggesting that these models increasingly extracted patterns found only in composite images and not representative of the real data set. These results are in good agreement with the general visual impression of style-transferred composite images being more representative of the real data. Most notably, the frequently observed dissonance between foreground and background in terms of saturation was eliminated during style transfer (Figure 3A-D).

**Figure 3.**
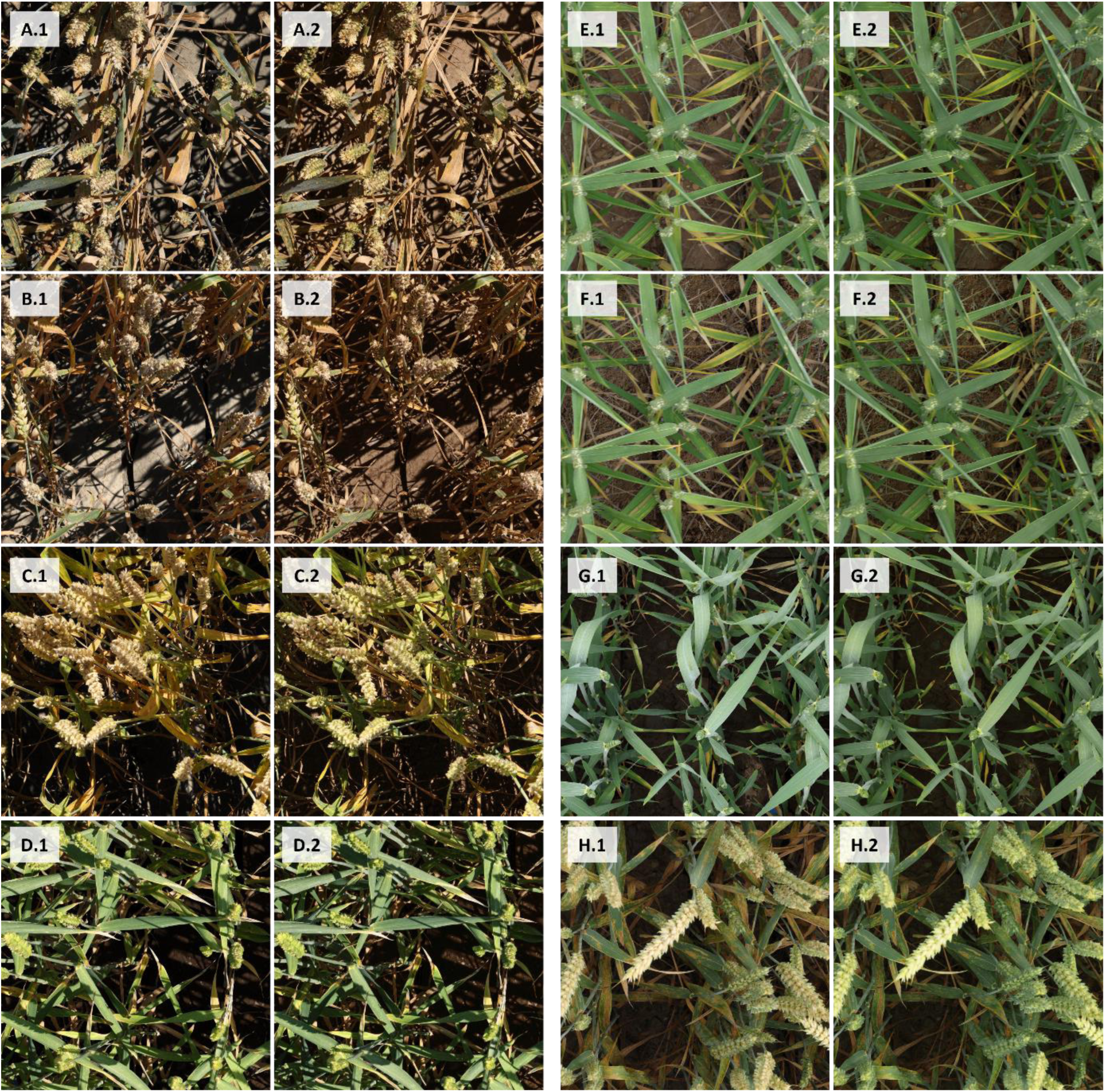
Composite images used for training of the vegetation segmentation models. Letters denote pairs of composite images before (raw; “.1”) and after style transfer (“.2”). **(A-D)** Images were captured under direct sunlight; **(E-F)** Images were captured under diffuse lighting. A separate CycleGAN was used for direct and diffuse lighting. (E) and (F) show identical plant foregrounds combined with different soil backgrounds.

Plant foregrounds could be combined with multiple randomly selected soil backgrounds (Figure 1F). Using a fixed CNN architecture and training procedure, training on ten instead of a single composite per plant foreground enabled an increase of the canopy segmentation model’s overall F1-score from 0.941 to 0.948, corresponding to a reduction of incorrect classifications by approximately 10% (Supplementary Figure S5).

### CNN-based semantic segmentation of high-resolution RGB images enabled the extraction of ear and shoot properties throughout the grain filling phase

Manual annotations of wheat ears from scratch on 180 image patches sized 1200 x 1200 pixels was sufficient to achieve useful performance also for the ear segmentation model, with an overall validation F1-Score of 0.89. A more detailed analysis of the performance metrics for both segmentation models at the level of individual validation images suggested that models performed equally well under direct and diffuse lighting as well as across phenological phases, i.e., during the stay-green and throughout the senescence phase (Supplementary Figures S6-S9). This allowed us to exclude a systematic bias from variable model performance across measurement dates. Some diverse examples illustrating the segmentation models’ performances are given in Figure 4.

**Figure 4.**
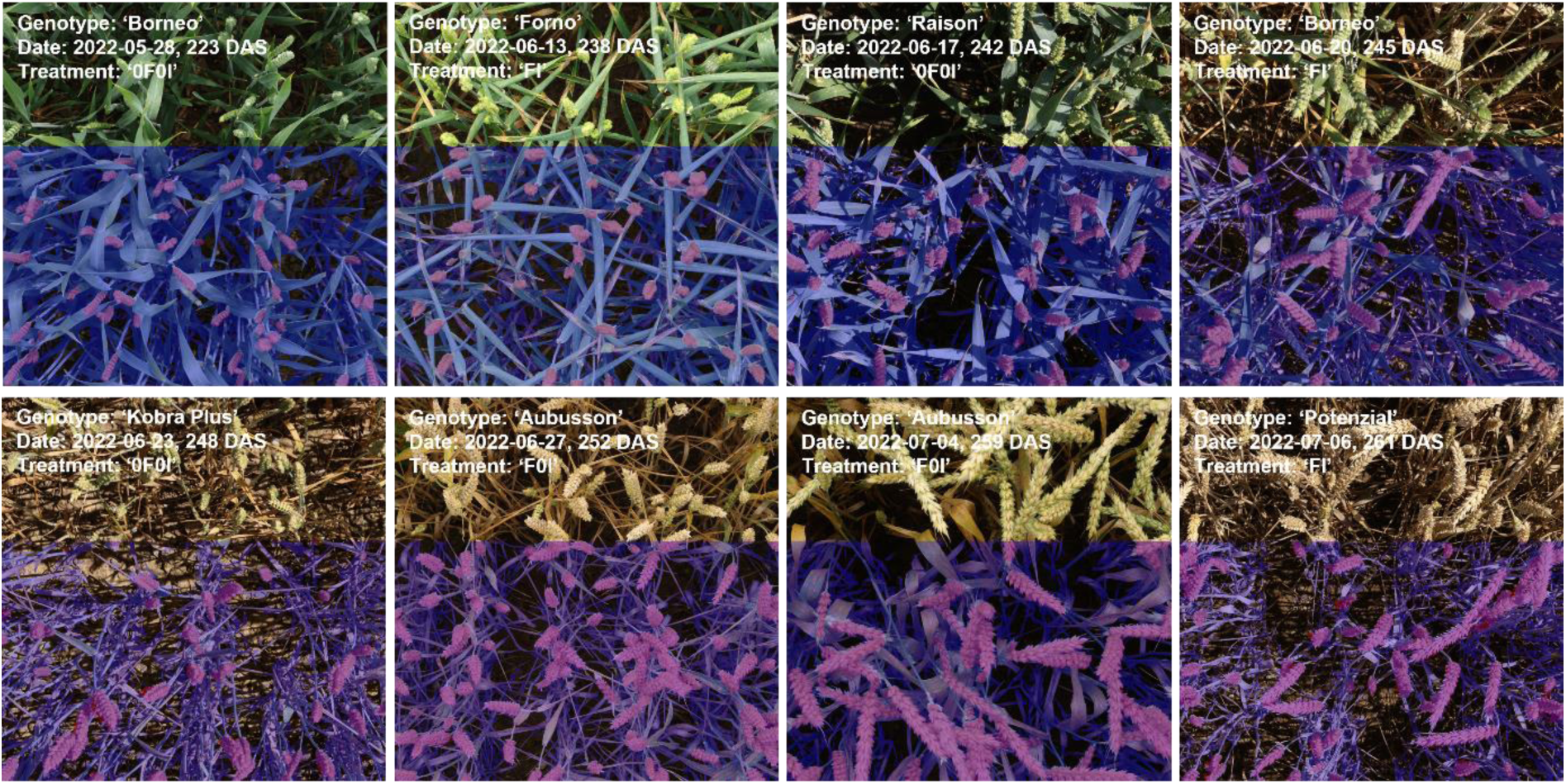
Inference on one randomly selected image for eight of the 17 measurement dates using the separate vegetation segmentation and ear segmentation models.

The cross-validated training accuracy (class-frequency-balanced) of the random forest classifier for classification of vegetation pixels into healthy/green, chlorotic/senescing, and necrotic/senescent tissue was 0.967 ± 0.006 (mean ± standard deviation across 10 folds). This high accuracy reflects the separability of the pixels in the training data set, particularly in HSV color space as well as some color indices (Supplementary Figure S10), which may in part be attributable to the somewhat biased training and validation data sampling strategy. However, training and validation data was sampled from numerous images representing strongly contrasting scenarios, and within those images from regions with strongly contrasting light exposure. Accordingly, the high accuracy also reflects the fact that, in comparison to the initial semantic segmentation, this 3-way classification is not a particularly challenging task (see Figure 5 for some examples). Interestingly, recursive feature elimination revealed that the RGB color space was not particularly useful for this classification. Instead, classifiers relying exclusively on the H channel of the HSV color space (H_HSV), u_Luv and the Excess greenness index (ExG) achieved near-optimal performance, with the addition of all other features having a negligible effect (Supplementary Figure S11).

**Figure 5.**
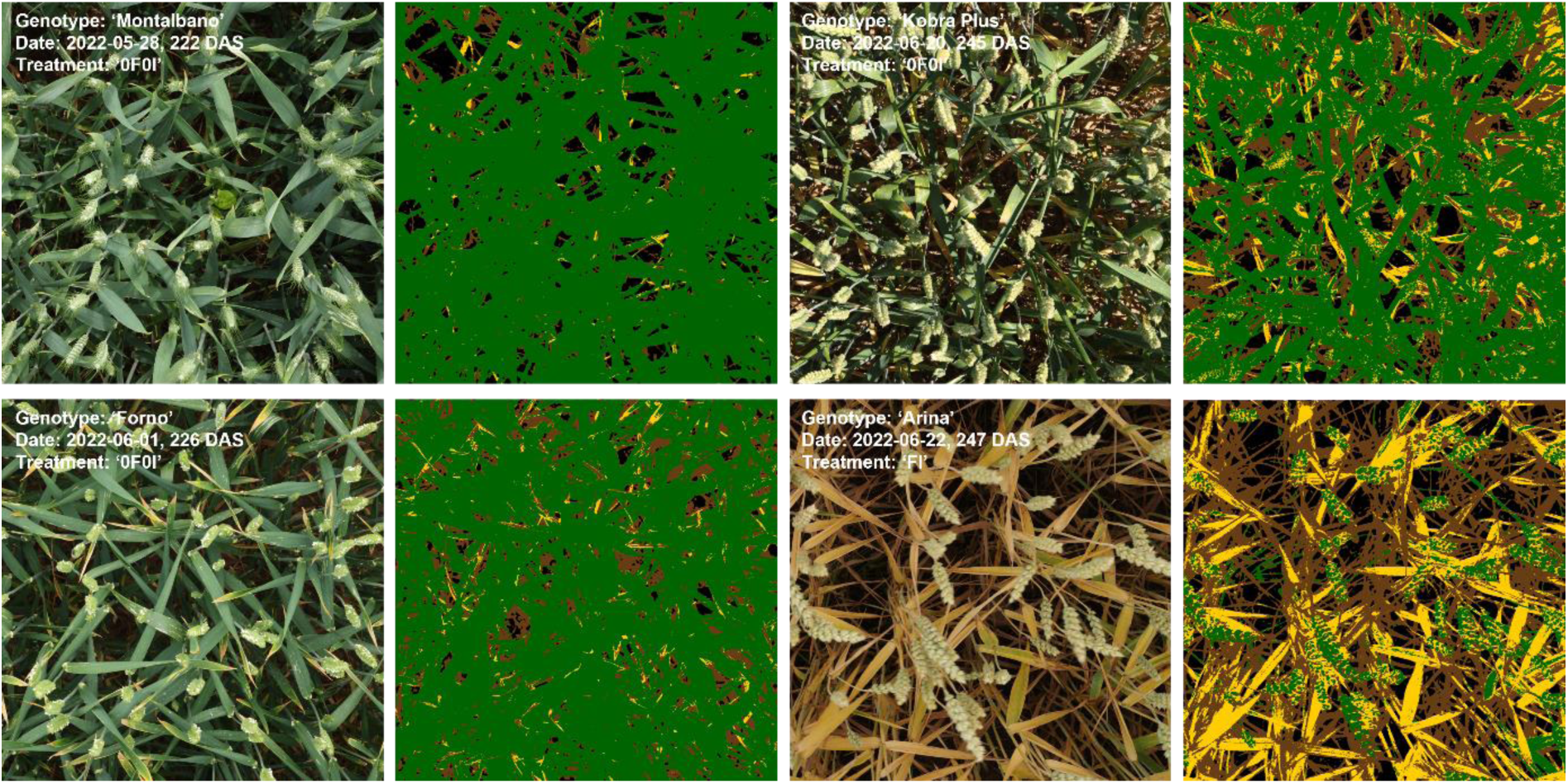
Prediction of vegetation status based on vegetation pixels color properties. Results are shown for one randomly selected image for four of the 17 measurement dates. Original images and corresponding prediction masks are shown, with green pixels indicating predicted healthy/green vegetation, yellow pixels indicating predicted senescing/chlorotic vegetation, brown pixels indicating predicted senescent/necrotic vegetation, and black pixels indicating predicted soil background.

### Segmentation of image time-series revealed dynamic patterns of vegetation cover and physiological status for ears and shoots

Plot-level time series of organ-level vegetation cover and vegetation status fractions further illustrate the high quality of the image segmentation. Both vegetation cover and vegetation status at the global and at the organ scale followed smooth temporal trends that were similar across all experimental plots and that could be very well explained (Figure 6, Figure 7, Supplementary Figures S12-S16). This is particularly noteworthy because our measurement setup did not guarantee that the exact same area of each plot was measured in subsequent images. Specifically, total vegetation cover showed a decreasing trend, whereas ear cover increased during grain filling in all plots. Consequently, shoot cover (i.e., vegetation cover without ears) showed a strongly decreasing trend, starting already approximately 10 d post-anthesis (Figure 6) and thus about two weeks earlier than the visually detected onset of canopy senescence in the earliest genotypes. The smooth temporal trends in all extracted traits clearly indicate a stable performance of the segmentation models irrespective of lighting conditions, genotype, treatment, or growth stage of the crop (Supplementary Figures S12-S14).

**Figure 6.**
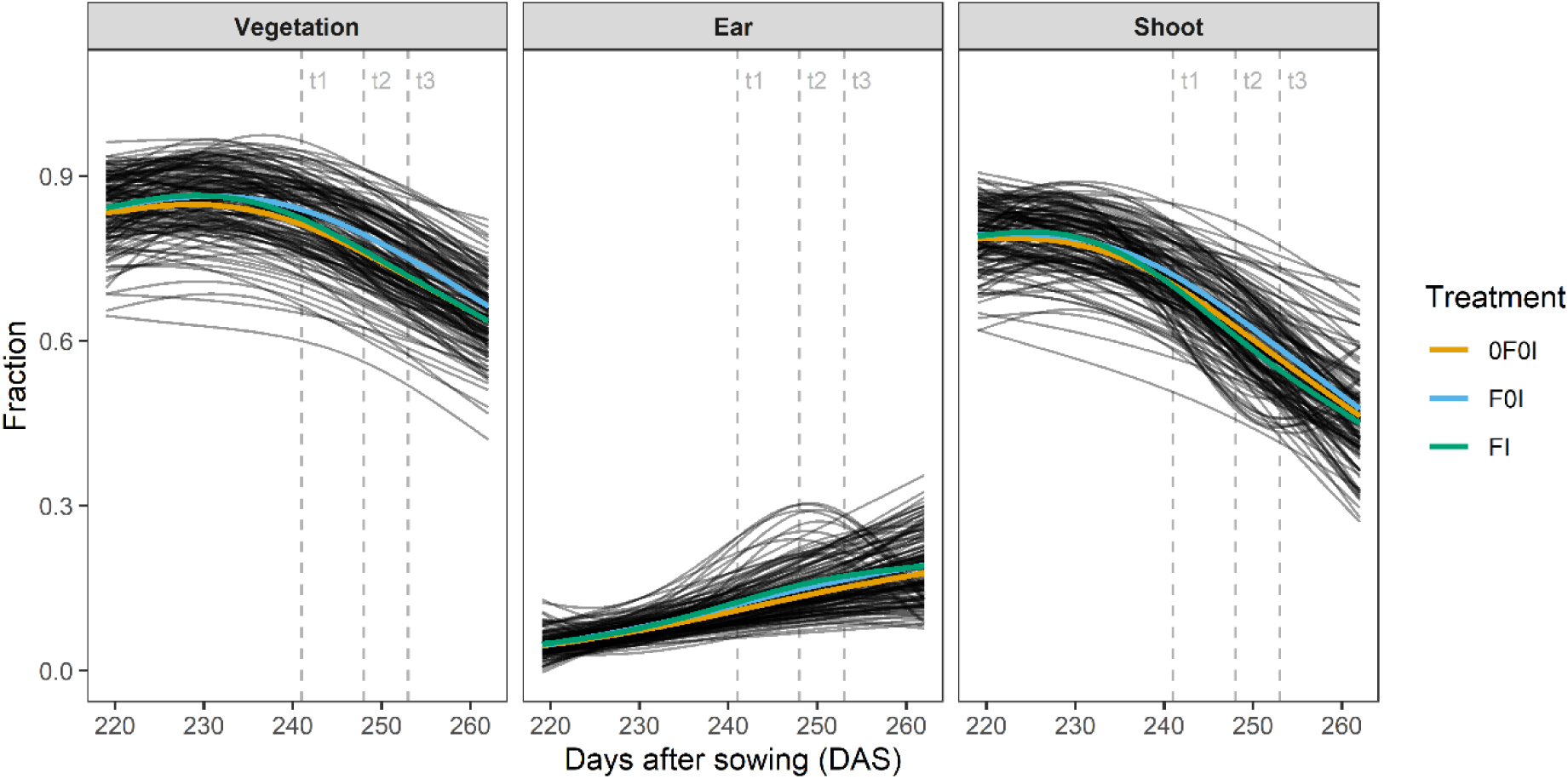
Fraction of images representing different components of vegetation, i.e., total vegetation, wheat ears, and vegetation without ears (i.e., leaves + stems = shoot) and their evolution over time between the first measurement at heading (May 25, 2022 [219 DAS, GS 55]) and the last measurement at physiological maturity (July 7, 2022 [262 DAS, GS 91]). Black curves represent p-spline fits to 17 data points for each experimental plot. Colored lines represent treatment means. Refer to Supplementary Figures for model fits and raw data at plot level.

**Figure 7.**
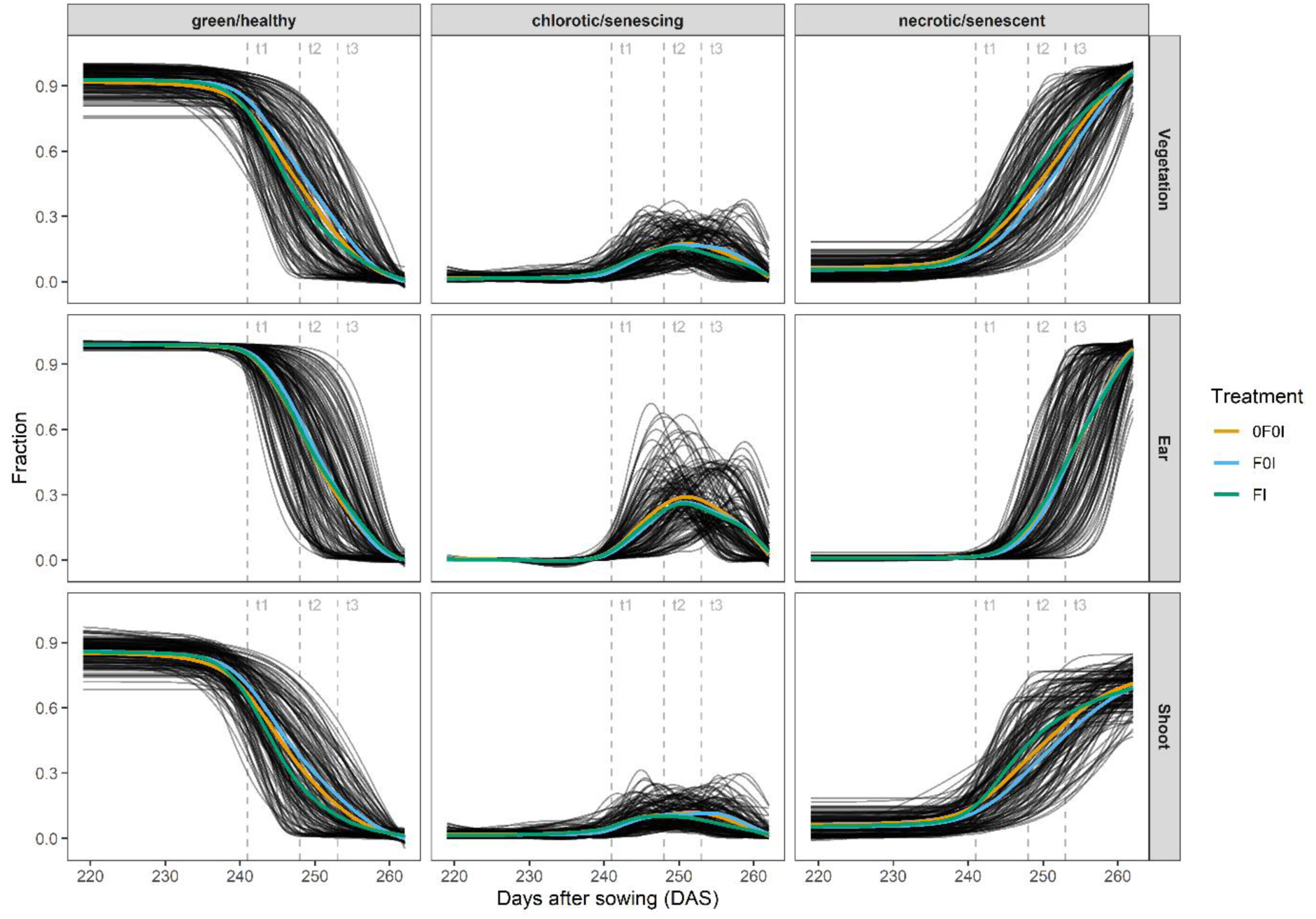
Relative contribution of healthy green, senescing/chlorotic, and senescent/necrotic tissue at organ level (total vegetation, shoot, and ears) and their evolution over time between the first measurement at heading (May 25, 2022 [219 DAS, GS 55]) and the last measurement at physiological maturity (July 7, 2022 [262 DAS, GS 91]). Black curves represent four-parameter Gompertz model fits or p-spline fits to 17 data points for each experimental plot. Colored lines represent treatment means. Refer to Supplementary Figures for model fits and raw data at plot level. Vertical dashed lines mark time points when the amount of STB was quantified.

The dynamics of the green fraction in shoots extracted from images closely followed the dynamics of the visual canopy greenness scores. Dynamics parameters extracted from image time series and from scorings were highly correlated, except for the onset (r = 0.41, r = 0.86, r = 0.91 and r = 0.87 for the onset, midpoint, end, and Integral, respectively; Figure 8). The trivial reason for the low correlation at the onset is the difficulty of defining an adequate threshold value across all plots, as it seems inappropriate to rescale fractions to a constant scale (Supplementary Figure S15). The fraction of the chlorotic/senescing tissue in vegetation components showed a peak during physiological senescence but was very low outside of this peak. This peak was clearly distinguishable in all plots and was more pronounced for ears than for shoots (Figure 7).

**Figure 8.**
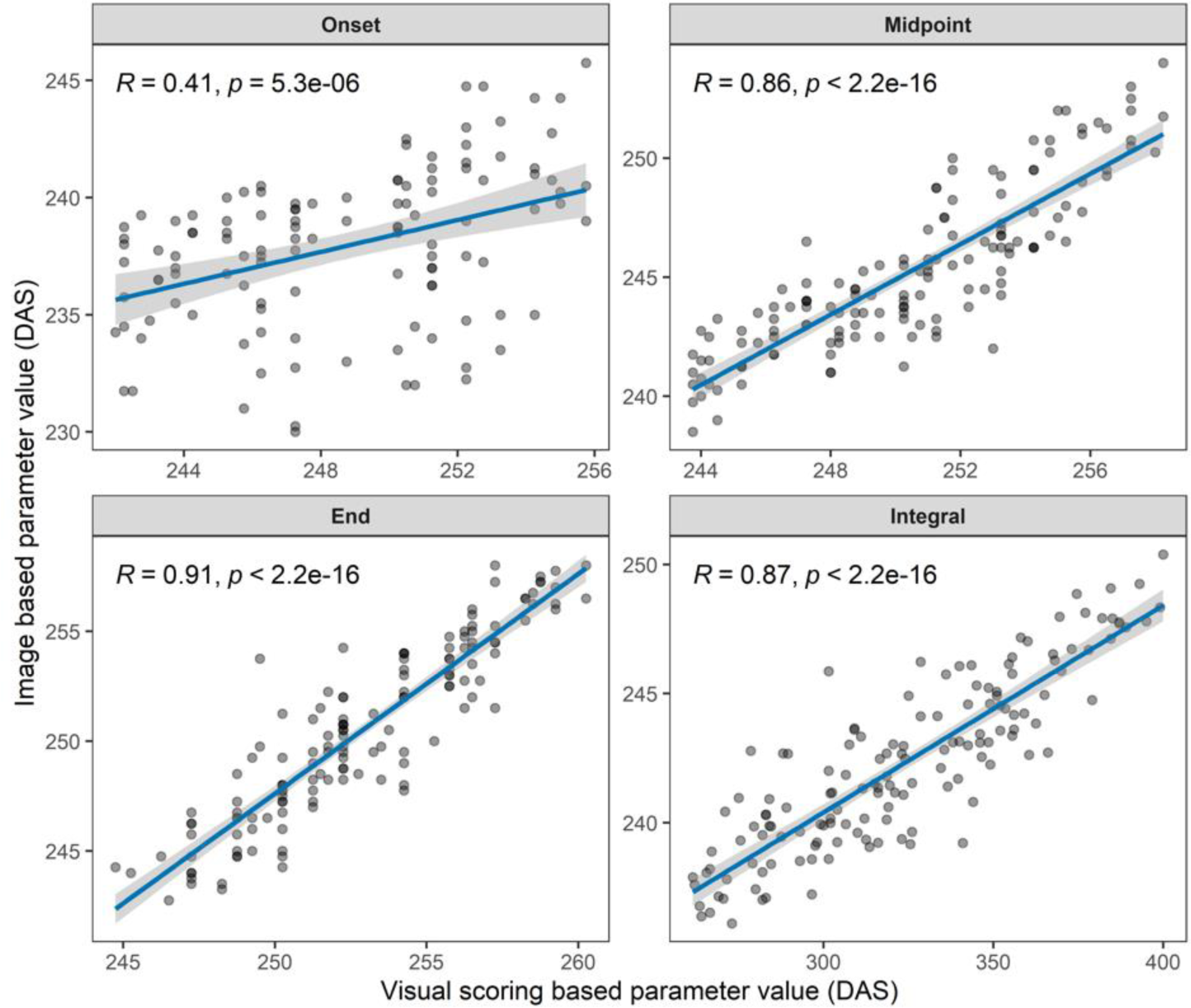
Pairwise correlation between canopy greenness decay dynamics parameters extracted from image time series and from visual scorings. Time points are reported in days after sowing (DAS). Pearson product moment correlation coefficients and p-values of the linear correlation are reported. The blue line represents the least squares line, the shaded ribbon represents the 95% confidence interval of the least squares line.

### Experimental treatments and genotype selection created a large variability in incidence and severity of foliar diseases

When averaging across all genotypes, there was a clear effect of the artificial inoculations on STB severity starting from t2 (Figure 9). The difference between inoculated and non-inoculated plots was primarily attributable to a strongly increased STB incidence, whereas conditional STB severity was more similar across treatments (Supplementary Figure S17, Supplementary Figure S18). In contrast, no strong effect of the inoculations was observable yet at t1, consistent with the long latency period of the disease of approximately 4 weeks. The development of the STB epidemic resulting from the artificial inoculations coincided with the onset of physiological senescence in many plots (Figure 7). Specifically, STB severity was still low at t1 even on flag leaf-1 (Figure 9), when first plots already showed signs of physiological senescence as indicated by increasing chlorosis and necrosis (Figure 7, Supplementary Figure S3). This was primarily the result of a generally early onset of senescence, which occurred about 8 d earlier than in a neighboring wheat experiment sown on the same date. Additionally, there were very strong spatial effects in the timing of senescence which accounted for more than 6 d differences (Supplementary Figure S19).

**Figure 9.**
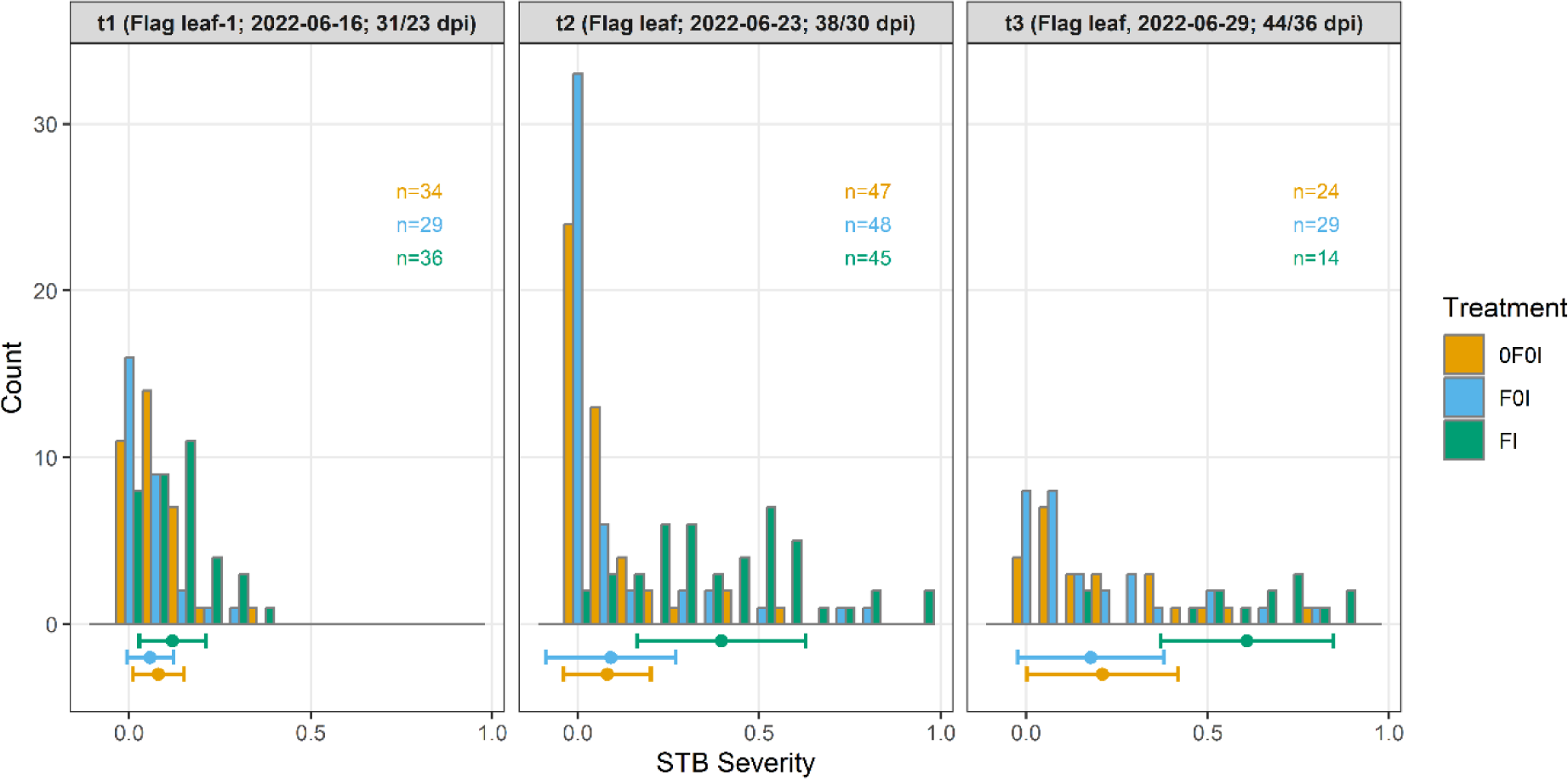
Distribution of septoria tritici blotch (STB) severity measured on three assessment dates (t1 – t3) during grain filling, depending on the treatments applied. STB severity was assessed as the product of visually scored disease incidence and image-based estimation of the percentage leaf area covered by STB lesions on infected leaves. At t1, assessments were made on a subset of plots where incidence was greater than or equal to 1/3, on flag leaf minus one. At t2 and t3, assessments were made on flag leaves. Points and horizontal whiskers below the bar plot represent per-treatment mean values and their standard deviations. Plot-based raw values are shown.

In terms of STB, there was no difference at the treatment level between the F0I and 0F0I treatments. This indicated that there was natural STB infection in these treatments which was not affected significantly by the early application of fungicide at GS 31. This is also supported by incidence assessments at t1 that revealed significant presence of STB infections especially at the flag leaf-1 and flag leaf-2 layers (Supplementary Figure S17). In contrast, the presence and severity of other foliar diseases, notably of yellow rust and brown rust, was strongly affected by the early application of fungicide (not shown). These diseases were clearly dominating in untreated plots, whereas they were virtually absent from fungicide-treated plots.

Besides the strong treatment effect, there was also ample variation of STB severity within treatments (Figure 9), largely attributable to genotypic differences in disease resistance and susceptibility. Genotypic means for STB severity based on individual, not spatially corrected plot values, at t2 ranged from 0.02 to 0.69, from 0.03 to 0.73, and from 0.10 to 0.84 for the treatments 0F0I, F0I, and FI, respectively.

### STB severity correlated with image-derived greenness decay dynamics and temporal patterns of chlorosis

The near-complete temporal overlap between the onset of the STB epidemic and physiological senescence (Figure 7) as well as strong spatial heterogeneities (Supplementary Figure S19) complicated the analysis of the effects of these traits on vegetation dynamics as reported in Figure 6 and Figure 7. Despite this, an overall treatment effect was observable on the dynamics of the green/healthy and the necrotic/senescent fractions of vegetation, particularly for shoots (Figure 7). Specifically, artificial inoculations (‘FI’ treatment) caused an earlier and faster decline in the green fraction of shoots and a concomitant earlier and faster increase in the necrotic/senescent fraction, when averaging across all plots within a treatment (Figure 7). A similar shift did not occur for ears. At the treatment level, no effects were observable on the temporal patterns of chlorosis.

When comparing treatment contrasts at the genotype-level based on spatially corrected plot values, a negative correlation between STB severity and the onset of canopy greenness decline was observed (Figure 10) suggesting that the developing STB epidemic was detected as an earlier decline in overall canopy greenness and confirming the above observation at the treatment level.

**Figure 10.**
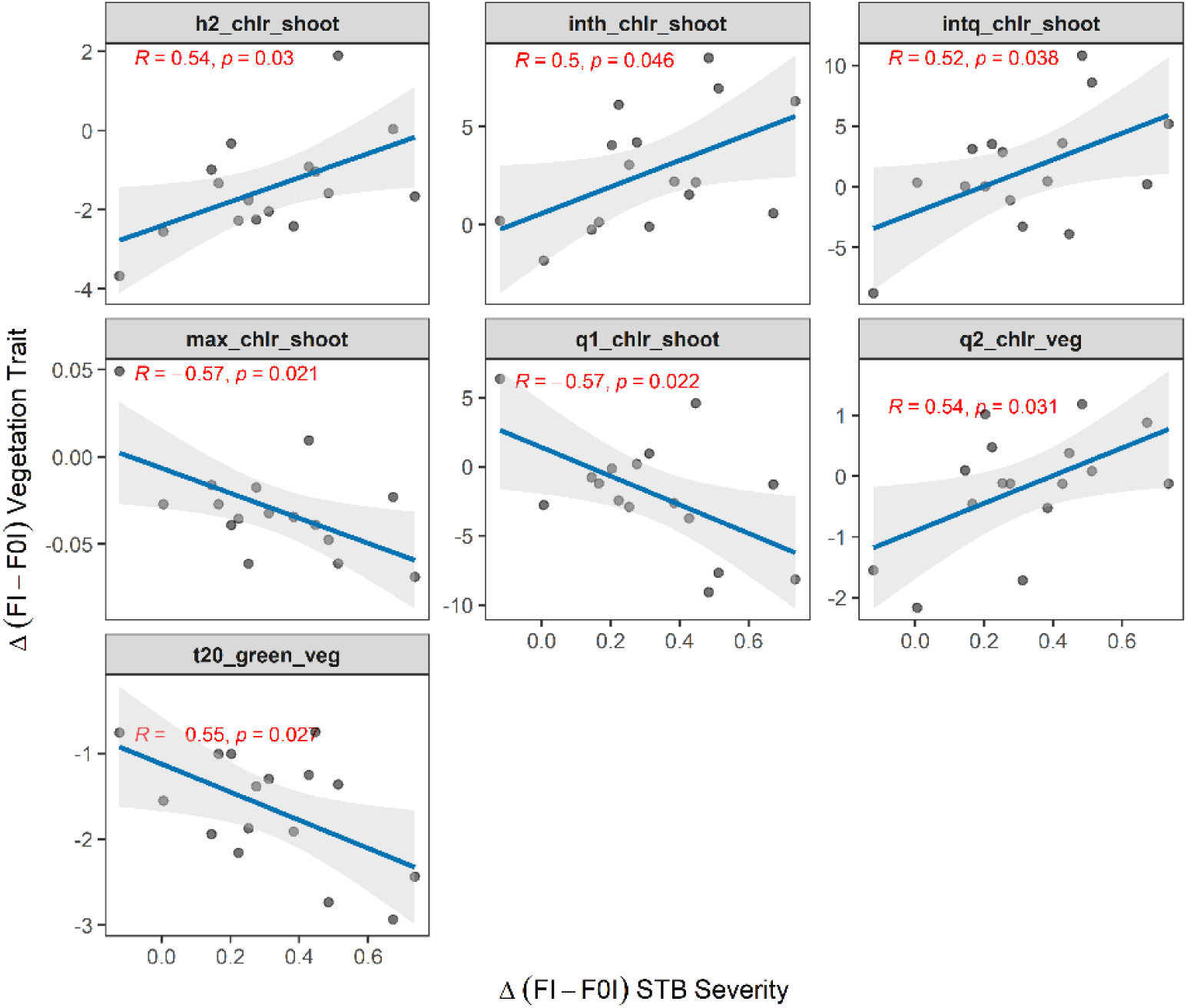
Pairwise correlation between the difference in STB severity and in vegetation fraction dynamics at the genotype-level (n = 16) as observed between the treatments ‘FI’ (early fungicide application + later inoculation – “clean STB”) and ‘F0I’ (early fungicide application, no inoculation – “healthy control”). Trait contrasts between treatments at the genotype-level were calculated as the difference of the mean corrected plot values for each genotype in each treatment. The parameters q1, q2, h1, and h2 denote the time points when one quarter and one half of the maximum was reached in the increasing and the decaying phase; the parameters intq and inth represent the durations q2 – q1 and h2 – h1, respectively (refer to Figure 2 for a graphical representation).

Besides the dynamics observed for the green and still healthy fraction, treatment contrasts in temporal patterns of chlorosis were also correlated with treatment contrasts in STB, which was not observed at the treatment level. A higher STB severity tended to result in a longer period during which significant chlorosis could be detected, but at the same time reduced the maximum chlorotic fraction detected in shoots (Figure 10). This suggested that, although no correlations between STB severity and the green fraction was observed during later stages, the relative contribution and timing of the necrotic and chlorotic fraction to the reduction in the green fraction differed depending on STB severity. Correlations for all extracted dynamics parameters are reported in Supplementary Table S2.

## Discussion

The main objective of this study was to develop methods enabling a detailed dynamic assessment of the physiological status of ears and shoots in wheat canopies throughout grain filling, as well as to develop perspectives on how such information may help elucidate the impact of stresses and stress responses. The following sections will therefore discuss the potential and limitations of the proposed methods and examine potential applications in the context of breeding for increased resistance and/or tolerance to biotic stress as well as optimized senescence dynamics.

### An efficient method to generate high-quality training data for vegetation segmentation under challenging conditions

In contrast to the annotation of readily recognizable individual organs, annotation of all vegetation in high-resolution images is challenging and extremely time-consuming. This is true particularly for maturing wheat canopies which are characterized by fine structures, very similar color properties as the soil background, and a complex canopy architecture resulting in complex lighting and shading patterns. Though a subsequent partitioning of the vegetation fraction to the organ-level is also challenging, individual organs such as ears are easier to recognize and, provided they stand out from their background, even weak bounding box annotations may provide a sufficient basis for the development of useful segmentation models (Dandrifosse et al., 2022b).

To overcome the limitation of generating sufficient training data for vegetation segmentation, we tested an approach based on image composition and domain transfer. Several earlier studies have implemented similar strategies or components of the strategy implemented here, especially in the context of weed detection or segmentation (e.g., Di Cicco et al., 2017; Fawakherji et al., 2020; Gao et al., 2020; Sapkota et al., 2022), although these studies typically had a strong focus on the generation of scenes with sparse vegetation and weed plants with a rosette-like growth habit. The approach presented here was chosen because it was expected to have high chances of success, since composite images contain all the features that are typically observed in real images of wheat stands, including all components of a stay-green or senescing wheat canopy, complex lighting, and richly textured and highly variable soil background. In contrast to other studies (e.g., Gao et al., 2020; Sapkota et al., 2022), the isolation of foreground instances was more challenging in our case due to the need to segment senescent plant tissues, and some manual annotation could not be avoided (Figure 1E). Nevertheless, the annotation effort was reduced about 20-fold with respect to annotations from scratch, and the risk for annotation errors was minimized. Data augmentation procedures - especially the application of color jittering - did not improve the performance of the segmentation model (Table 1), indicating that the synthetic training images were highly representative of the real-world data set. Visual examination of the domain-transferred composite images occasionally revealed atypical features such as small patches of greenish soil (see e.g., Figure 3E, 3F), especially in cases where the original soil background image had low saturation. It may therefore be that the domain transfer introduced some noise that resulted in a more diverse training data set than could have been obtained through manual annotation of a set of images from the target domain.

The datasets generated in the context of this study explicitly cover a wide range of scenarios in terms of genotype morphology, healthiness, canopy structure, soil background, lighting conditions, and phenology. In contrast, robustness of trained segmentation models to variation in imaging set-up, type of vegetation, or camera parameters was not addressed here and may limit the usefulness of the data sets in other contexts, as compared to the data set described by Serouart et al. (2022). However, our approach is directly applicable to additional scenarios (future wheat experiments, or even experiments involving other crops), which facilitates an expansion of the data set with low effort.

Our results encourage more detailed studies that should evaluate whether fully synthetic training data could be obtained, e.g., through a combination of functional-structural plant growth models such as ADEL-Wheat (Fournier et al., 2003) with open-source rendering software such as Blender (Blender foundation, https://www.blender.org). Promising results have been achieved using similar approaches in other contexts, e.g. for Arabidopsis (Ubbens et al., 2018) and sweet pepper (Barth et al., 2018).

### Color-based inference of vegetation status at organ-scale using high-resolution RGB imagery

The identification of the vegetation fraction and the separation of ears and shoots enabled a component-level analysis of color properties in this study. The three-way classification done here represents one amongst several possible ways to gain insights into the relative contribution of different vegetation fractions. A similar approach was used by Makanza et al. (2018) who classified pixels in aerial images into ‘green’, ‘yellow’ and ‘dry brown’, interpreting these classes as representing differentially advanced stages of canopy senescence. Serouart et al. (2022) performed a two-way classification of previously segmented vegetation pixels into green and senescent pixels, considering both chlorotic and necrotic vegetation as ‘senescent’. In agreement with our results, this study also highlighted the need for color space conversion in order to achieve a robust classification (Serouart et al., 2022). Other studies used color indices to infer vegetation status or separate green from senescent vegetation (Anderegg et al., 2023; Rasmussen et al., 2019; Schirrmann et al., 2016). Here, it was reasoned that chlorosis and necrosis represent readily visually distinguishable vegetation fractions with a distinct physiological status. Widespread chlorosis is typically indicative of a tightly controlled senescence process encompassing a degradation of chloroplasts as well as major changes in the chlorophyll/carotenoid ratio resulting from differential breakdown rates of these pigments during early senescence (Fischer and Feller, 1994; Lim et al., 2007; Sanger, 1971). The involved controlled degradation processes support nutrient remobilization and subsequent nutrient translocation to developing grains, which is critical for yield and quality formation in wheat (Kichey et al., 2007). In contrast, direct necrosis causes a complete loss of function and the trapping of resources in the affected tissues. Therefore, chlorotic and necrotic fractions should be considered separately. The color-based approach commonly used to achieve this separation may be limited in precision, because (i) the senescence-related transition from green to chlorotic and ultimately necrotic tissue is a continuous process, meaning that a definition of thresholds is inherently arbitrary and subjective, and (ii) reliance on the visible spectrum may be insufficient to precisely detect the switch from stay-green to senescence (Jagadish et al., 2015; Merzlyak et al., 1999). However, even if some degree of subjectivity remains, the application of well-defined decision boundaries irrespective of the context should guarantee unbiased estimates of vegetation fractions across trials and genotypes. Furthermore, with a view towards high throughput applicability under field conditions, spatial resolution and thus the possibility to eliminate background signal and extract properties separately for different vegetation components may offer significantly larger benefits than a high spectral resolution, because vegetation cover and the relative contributions of different vegetation components change massively during the final growth stages (Figure 6). Such changes are highly likely to be genotype-specific, thus introducing a large bias in canopy-level signals if disregarded (Anderegg et al., 2020). In the future, merging different types of sensor data to extract component-level information beyond the visible spectral domain may offer significant potential for more precise and objective measurements (Dandrifosse et al., 2022a; Jagadish et al., 2015).

### Leveraging vegetation dynamics to track epidemics and separate effects of foliar diseases and physiological senescence on canopy greenness decay

Repeated imaging enabled the extraction of temporal patterns in vegetation fractions for ears and shoots during the development of the STB epidemic and senescence using dynamic modelling. Despite the challenging outdoor conditions varying strongly across imaging time points (see Figure 4), our segmentation pipeline enabled the extraction of clear temporal patterns from repeated imaging (Figure 2, Figure 7, Supplementary Figures S15 and S16). The observed patterns are in good agreement with the expectation that the green fraction should show a monotonous decrease that can be well described using parametric models (Figure 2, Figure 7, Supplementary Figure S15; Anderegg et al., 2020; Bogard et al., 2011; Christopher et al., 2014). The chlorotic fraction showed a peak during rapid physiological senescence but was virtually absent outside of this restricted time window, which is well in line with the interpretation of chlorosis as a phenotypic marker for physiological senescence.

The ability to track the dynamic development of the green, chlorotic and necrotic fractions at component level offers new opportunities to assess the impact of stresses as well as stress responses. First, repeated assessments of the green, chlorotic and necrotic fractions may enable a precise detection of the moment when leaf senescence or leaf disorders start to affect the light absorption capacity of the upper-most leaf layers, irrespective of changes in organ or background contribution to the scene. While both foliar diseases and physiological senescence ultimately lead to widespread necrosis, the sequential development of the symptoms differs markedly: the development of necrotic lesions caused by pathogens is not typically preceded by chlorosis to a similar extent. Necrosis remains restricted to scattered regions on leaves, thus causing necrotic islands in green tissues. In contrast, widespread chlorosis should mark the onset of physiological senescence. Consequently, a separate quantification of the chlorotic and necrotic vegetation fractions should provide insights into the drivers of greenness decay, where a large contribution of chlorosis (both in terms of the fraction of vegetation affected and in terms of the duration of its persistence) indicates a strong contribution of physiological senescence, whereas prominent necrosis without a gradual transition through chlorosis indicates leaf damage. These leaf damages are likely to result from biotic stress, because abiotic stress is well known to accelerate physiological senescence (Bogard et al., 2011; Distelfeld et al., 2014; Martre et al., 2006), but does not typically cause localized leaf damage.

Despite the near-simultaneous onset of the STB epidemic and physiological senescence and pronounced field heterogeneity, our analysis revealed an effect of STB especially on the dynamics and prominence of chlorosis in shoots (i.e., the disease-affected fraction; Figure 10). Specifically, higher STB severity coincided with a lower contribution of chlorosis to the overall greenness decay, in line with our above interpretations. A detailed comparison of the temporal dynamics of chlorosis and necrosis is likely to reveal potential and currently elusive interactions between disease-induced necrosis and physiological senescence. This is highly relevant, since the capacity of genotypes to maintain the functionality of remaining healthy leaf area and avoid an anticipation of (or even delay) the onset of physiological senescence in the presence of disease may represent an important compensation mechanism leading to tolerance. Anticipating the onset of the STB epidemic with respect to the onset of physiological senescence (e.g., through earlier inoculations) will allow for more detailed investigations of the interactions between foliar diseases and physiological senescence. The potentially high-throughput nature of the proposed methods paves the road to genetic studies in this direction.

Finally, an interesting observation was that ear senescence patterns appeared to be unaffected by the presence and severity of STB. In contrast, we frequently observed an extended persistence of green stems in heavily diseased inoculated plots. We hypothesize that this may represent a response to premature losses in green leaf area, facilitating a more complete remobilization of stem reserves to sustain concurrent grain filling. Further distinguishing between stems and leaves within the shoot may therefore reveal additional compensation mechanisms using the same data sets. Additionally, the separation of stems including peduncles would greatly enhance the scoring of physiological senescence, as peduncle senescence is considered a reliable measure for this trait (Chapman et al., 2021).

### Limitations of the approach in terms of disease detection and quantification

Image data collected here has sufficient resolution to enable an easy recognition of individual disease symptoms such as necrotic lesions or rust pustules. However, a precise diagnosis especially for necrotrophic diseases requires an even higher resolution (Karisto et al., 2018; Stewart et al., 2016), because unique features attributable to certain diseases such as black fruiting bodies (pycnidia) within necrotic lesions in the case of STB may be microscopic. Methods for in-field disease detection and quantification in stay-green canopies using very high-resolution imagery are currently being developed (Zenkl et al., unpublished), and will represent an important complement to the methodologies developed in this study.

In terms of disease quantification, nadir imagery is always limited by the restricted visibility of lower leaf layers. This is particularly problematic in the case of STB as the symptoms are typically more visible on the lower leaves and the disease moves into the top leaf layers via splash-dispersed spores, followed by a long latent period. Hence an early detection of the disease using nadir images is practically impossible. Whereas early disease detection is not a primary objective in breeding, it is key for the implementation of concepts of precision agriculture. Successful early detection of STB will arguably always have to involve some sort of physical interaction with the crop to expose lower leaf layers. Here, we aimed to maximize the visibility of lower leaf layers by setting an optimal exposure bias within the tolerable range. High dynamic range cameras may offer interesting opportunities to minimize this issue, but occlusion will remain the dominant problem is this respect.

## Conclusions

The use of state-of-the-art deep learning models for image segmentation and a subsequent color-based classification and dynamic modelling facilitated a time-resolved monitoring of the physiological status of vegetation separately for ears and shoots. Application of these methods to image time series allowed for an accurate reproduction of visually observed greenness decay dynamics and revealed contrasting temporal patterns of greenness decay and chlorosis in plots differing with respect to their infestation levels with foliar diseases. The observed patterns are in good agreement with an interpretation of chlorosis as a phenotypic marker of physiological senescence, suggesting that a separate analysis of the chlorotic and necrotic vegetation fraction in disease-affected vegetation components may facilitate a separation of the effects of foliar diseases and physiological senescence on overall greenness dynamics. Thus, the developed tools hold significant potential for high throughput assessments of crop responses to biotic stress under field conditions.

## General

We thank Julien Alassimone (ETH Plant Pathology) for instructions and practical assistance in preparing the artificial pathogen inoculations. We gratefully acknowledge support from the Group of Crop Science at ETH Zürich, especially Simon Corrado for crop husbandry, Norbert Kirchgessner for technical advice regarding data acquisition, and Brigitta Herzog for assistance with seed preparation and management. Manual annotations were supported by Seraina Wagner, Giulia Malingamba, Sara Holdener, and Juliane Ebenhög. We also thank Christoph Grieder for managing the 2015 field experiment and Lukas Kronenberg for acquiring the image data for that experiment in the framework of his BSc thesis. Inspiration for the use of CycleGANs was taken from a student report crafted in the framework of the “Data Science Lab” at ETH Zürich by Fabian Bosshard, Matteo Guscetti, and Lukas Hächler, supervised by Lukas Roth and Prof. Ce Zhang.

## Author contributions

Jonas Anderegg: conceptualization, methodology, software, investigation, data curation, formal analysis, visualization, writing – original draft. Radek Zenkl: investigation, software, writing – review & editing. Achim Walter: resources, writing – review & editing. Andreas Hund: conceptualization, resources, writing – review & editing. Bruce McDonald: funding acquisition, resources, writing – review & editing.

## Data Availability

All code and data sets pertaining to deep learning model optimization and training will be open-sourced on Github, an archived version will be made available *via* the ETH Zürich publications and research data repository (https://www.research-collection.ethz.ch/). Additional raw data will be made available upon reasonable request.

## Footnotes

1 The python implementation of the algorithm from https://github.com/kuan-wang/The_Bilateral_Solver was used.

2 https://www.cvat.ai/

3 https://github.com/junyanz/pytorch-CycleGAN-and-pix2pix

