## Supplementary Materials for "Combining high-resolution imaging, deep learning, and dynamic modelling to separate disease and senescence in wheat canopies"

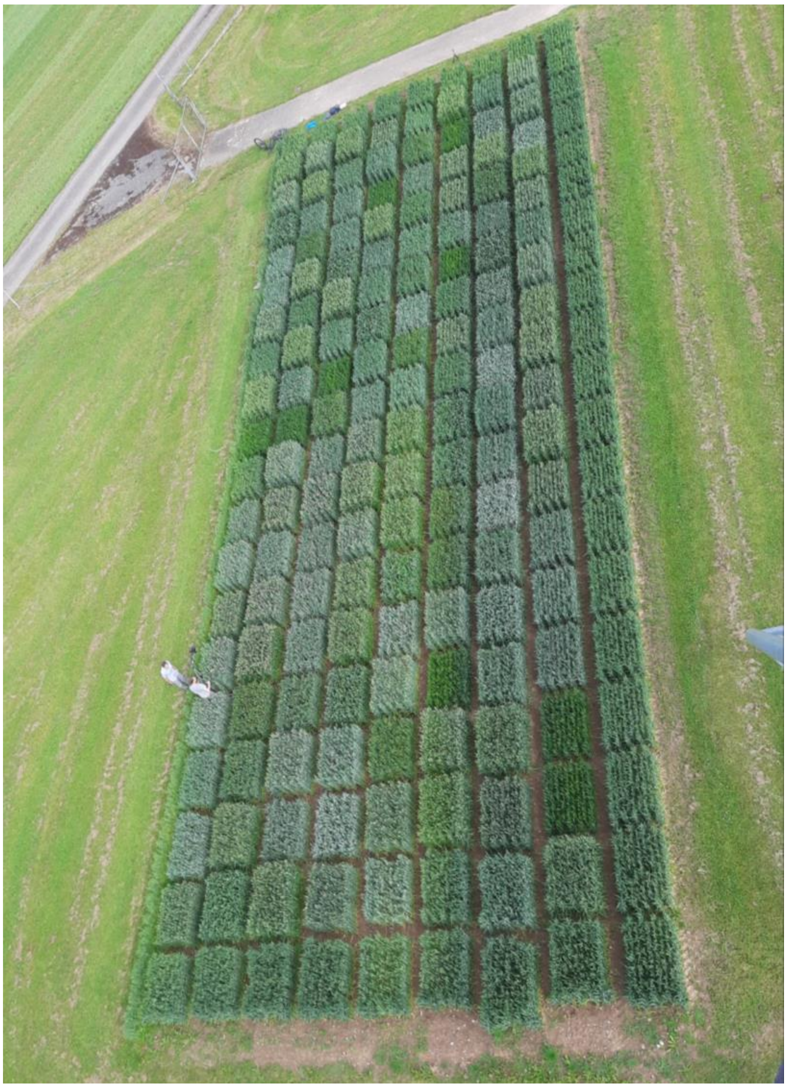

***Supplementary Figure S 1*** *Bird’s eye view of the field experiment. The selection of wheat genotypes for this experiment resulted in strongly varying physical appearance of wheat stands due to contrasting leaf glaucousness and flag leaf angle. Image courtesy Norbert Kirchgessner, Crop Science Group, ETH Zürich.*

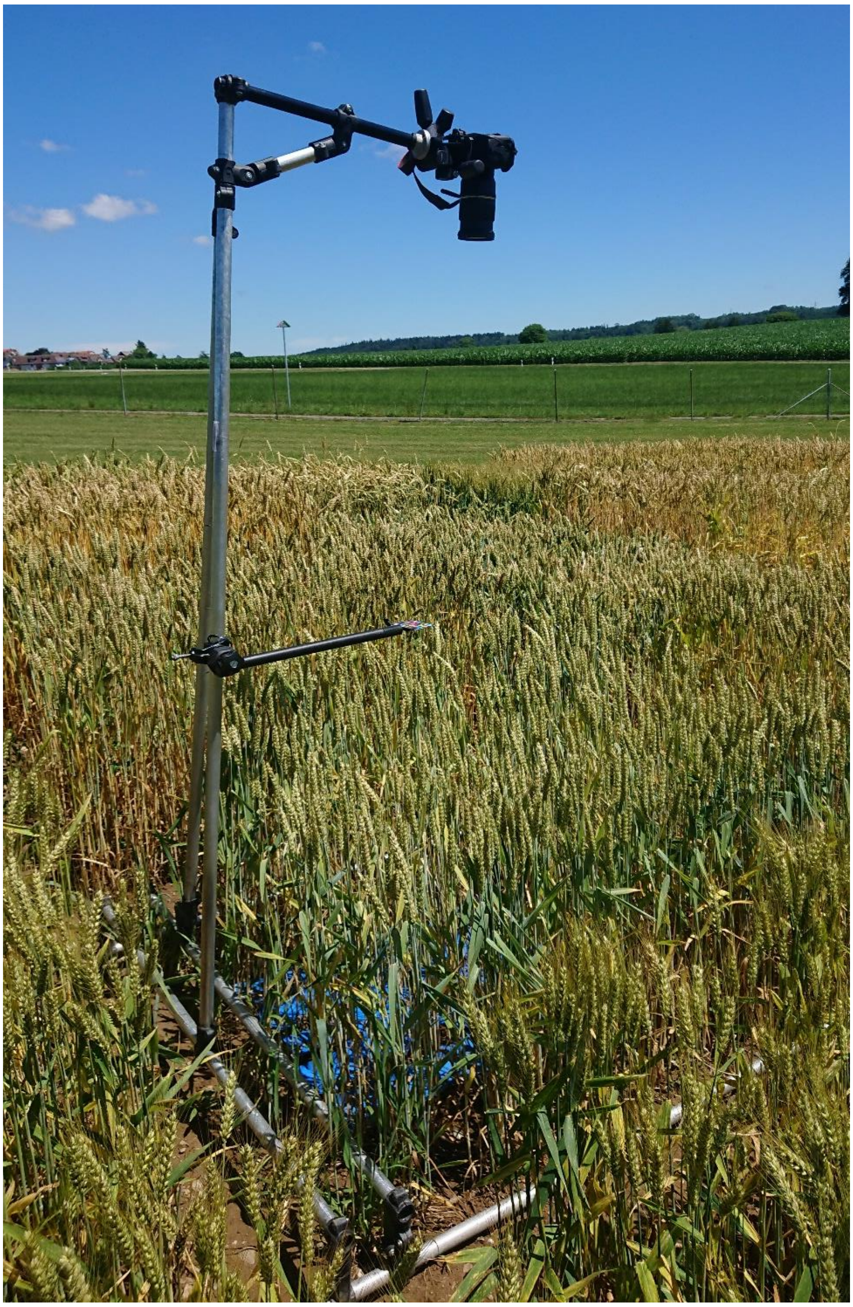

***Supplementary Figure S 2*** *Imaging setup.*

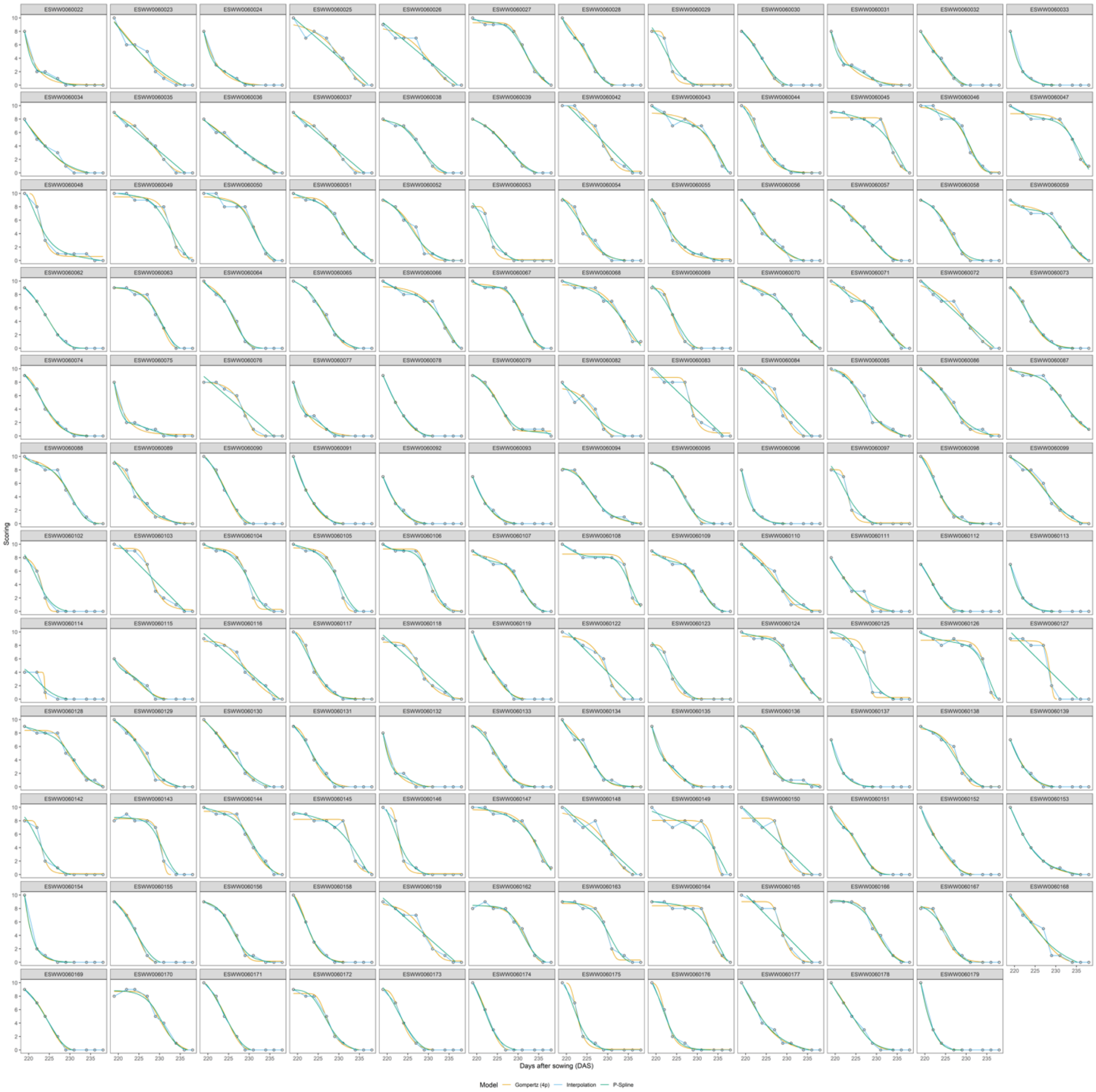


***Supplementary Figure S 3*** *Plot-level raw data for the fraction of healthy green leaf area in the total leaf area as estimated visually. Visual scorings on the y-axis are plotted against the time-point of assessment, indicated in days after sowing, on the x-axis. Each facet represents raw data for one experimental plot (black circles, nine scorings between the onset of senescence in the earliest genotype and physiological maturity in the latest genotype). Plot time series were fitted using a four-parameter Gompertz model, P-splines, and linear interpolation. Please zoom in for details.*

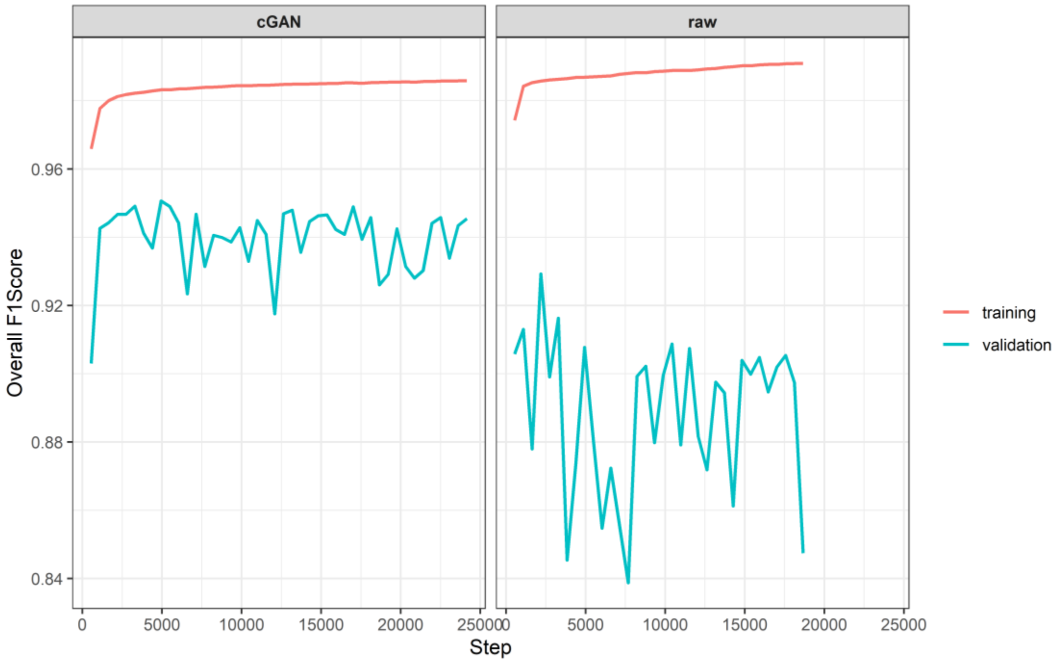


***Supplementary Figure S 4*** *Comparison of the performance of vegetation segmentation models trained on raw and on style-transferred composite images. Models were evaluated on real images (74 manually annotated patches sized 1200 x 1200 pixels). Training was stopped when no improvement in the overall validation F1-Score was achieved for 30 epochs of training.*

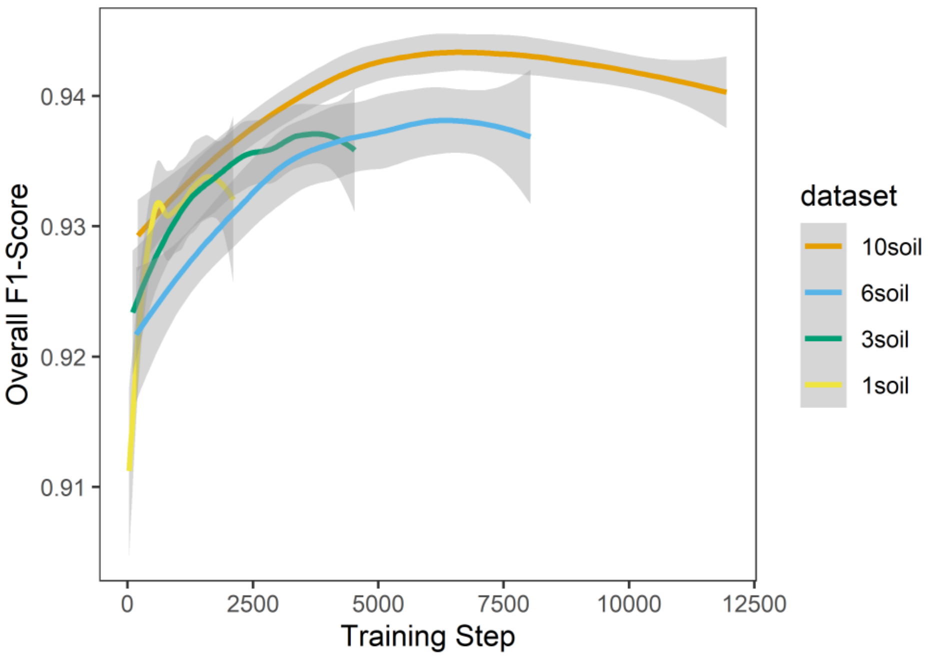


***Supplementary Figure S 5*** *Overall validation F1-Score of the vegetation segmentation model throughout the training process, depending on the training data set used. Different data sets consisted of plant vegetation foregrounds combined with 1, 3, 6 and 10 randomly selected soil backgrounds. Colored lines represent loess - smoothed values, shaded ribbons represent 95% confidence intervals. The training data set with 10 soil backgrounds represents the full data set, with the other datasets representing independent random subsets (i.e., datasets are not nested). Training was stopped when no improvement in the overall validation F1-Score was observed for 20 epochs.*

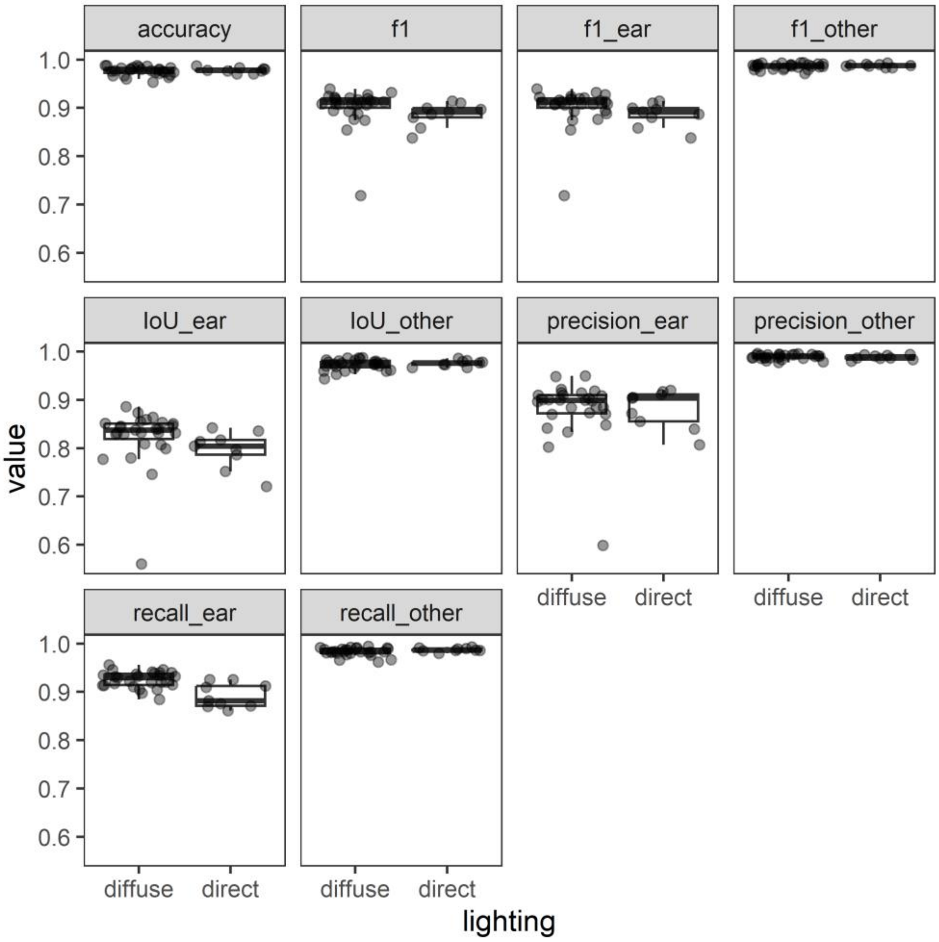


***Supplementary Figure S 6*** *Performance metrics for the ear segmentation model depending on the lighting conditions ('diffuse' and 'direct') present for the validation images (n = 36). Where meaningful, metrics are reported separately for the two classes considered by the model, i.e., 'ear' and 'other'.*

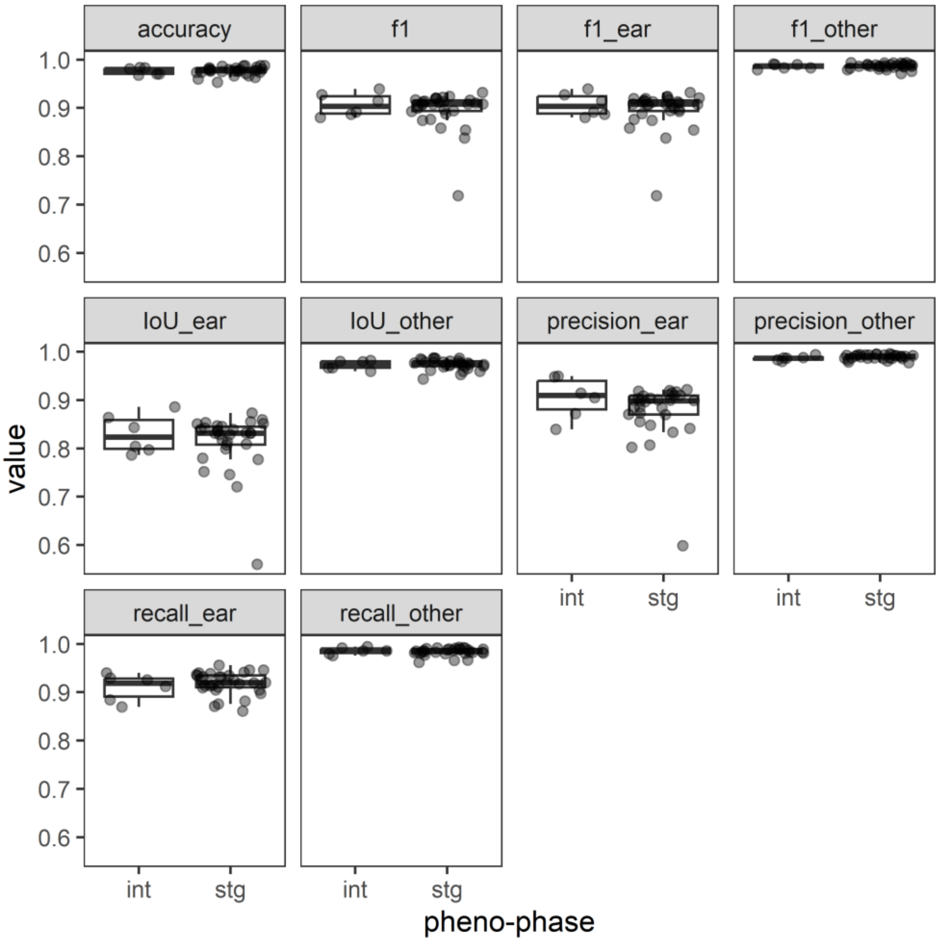


***Supplementary Figure S 7*** *Performance metrics for the ear segmentation model depending on the phenological phase represented by the validation images (n = 36). Validation images were classified into images representing stay-green canopies ('stg') and images representing canopies at an intermediate stage of senescence ('int'). Where meaningful, metrics are reported separately for the two classes considered by the model, i.e., 'ear' and 'other'.*

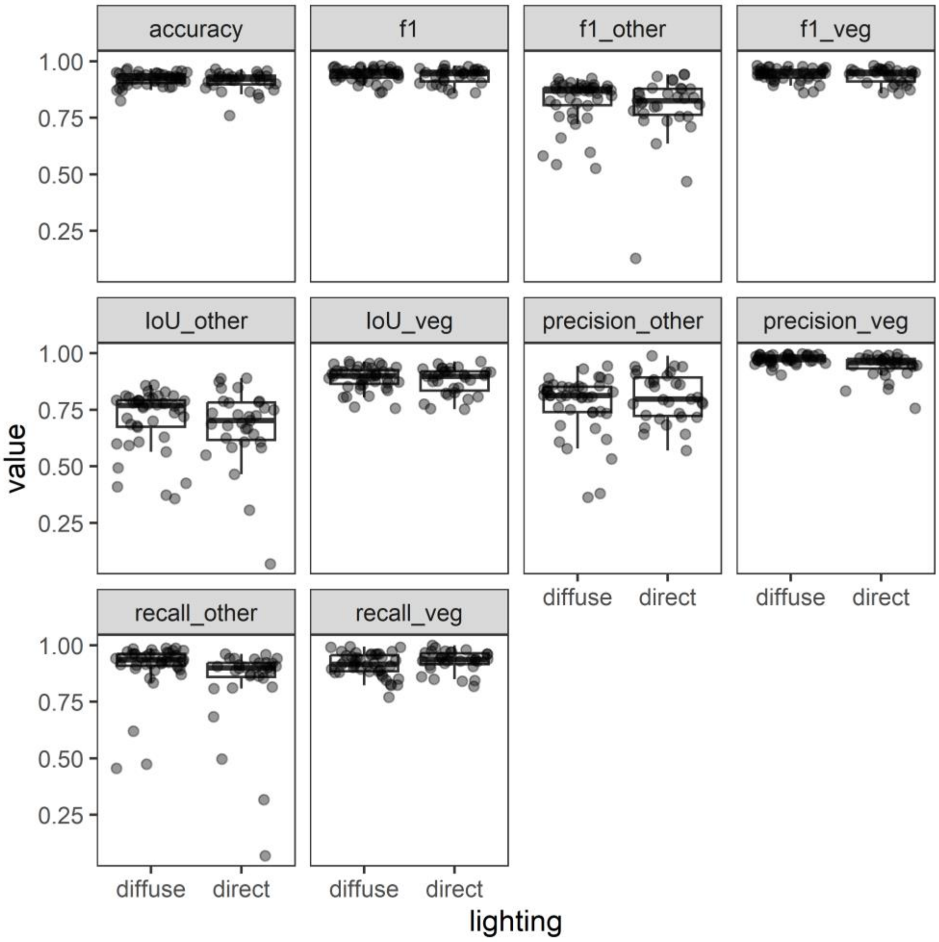


***Supplementary Figure S 8*** *Performance metrics for the vegetation segmentation model depending on the lighting conditions ('diffuse' and 'direct') present for the validation images (n = 74). Where meaningful, metrics are reported separately for the two classes considered by the model, i.e., 'vegetation' ('veg') and 'other'.*

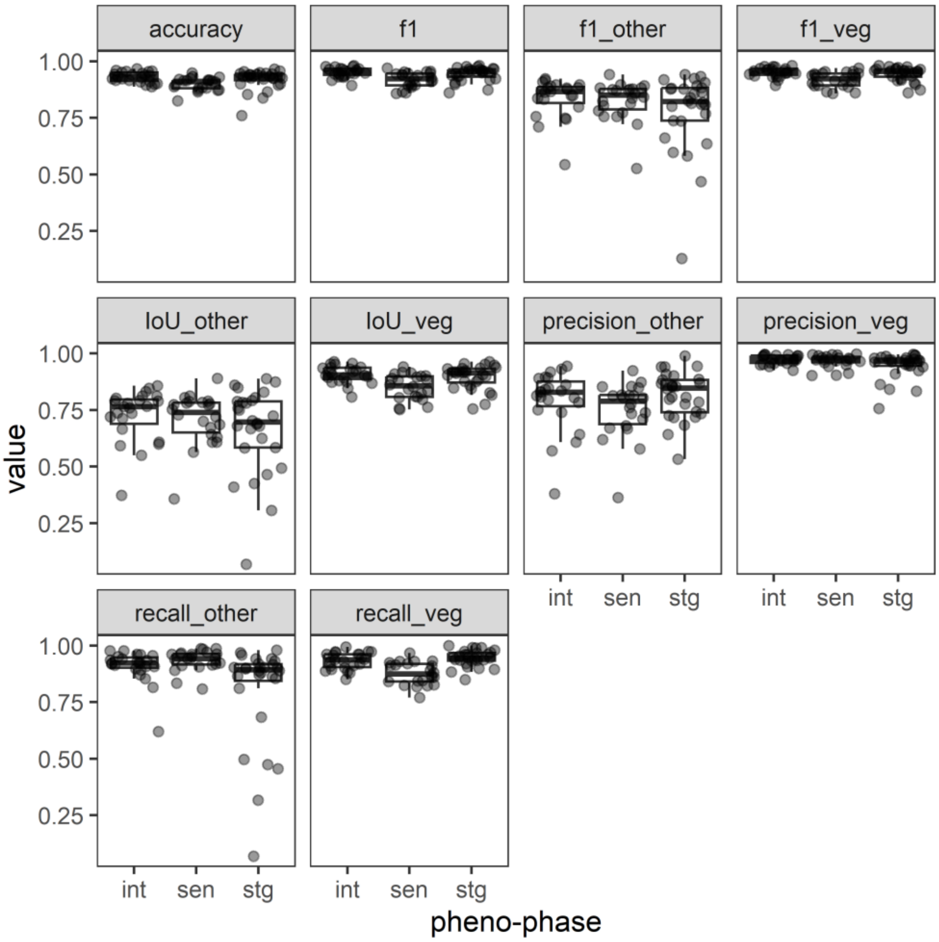


***Supplementary Figure S 9*** *Performance metrics for the vegetation segmentation model depending on the phenological phase represented by the validation images (n = 74). Validation images were classified into images representing stay-green canopies ('stg'), images representing canopies at an intermediate stage of senescence ('int'), and images representing canopies at an advanced stage of senescence ('sen'). Where meaningful, metrics are reported separately for the two classes considered by the model, i.e., 'vegetation' ('veg) and 'other'.*

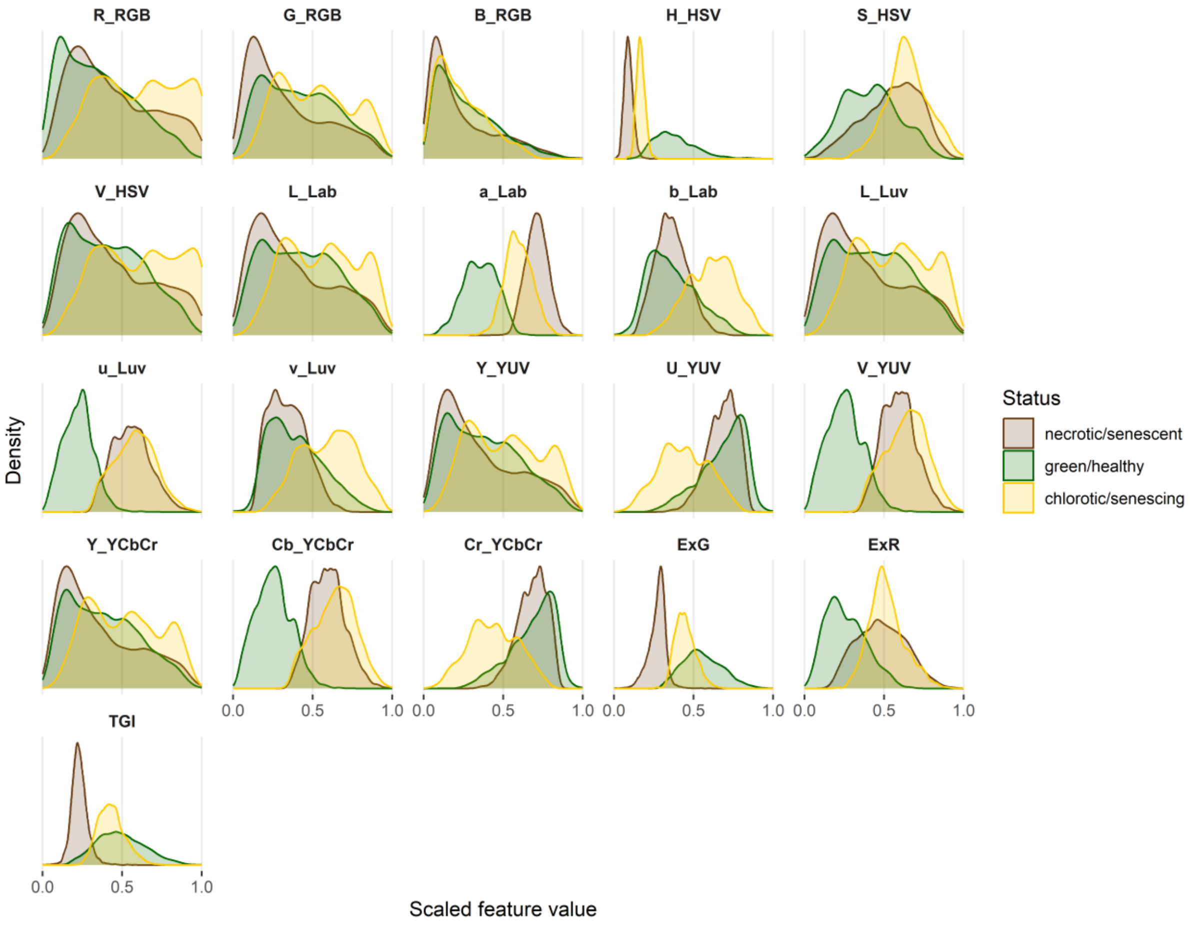


***Supplementary Figure S 10*** *Density plots illustrating the distribution of feature values in the sampled training and validation data for the three classes of vegetation. Feature values were scaled to the range [0; 1]. These color features served as predictors to assign vegetation pixels to one of the three classes of vegetation.*

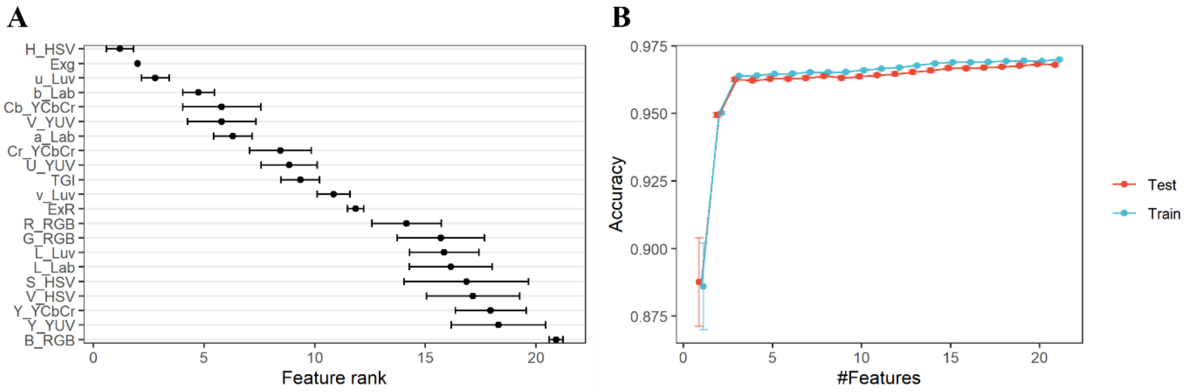


***Supplementary Figure S 11*** *Feature selection for prediction of the vegetation class using pixel color information.* ***(A)*** *Feature importance ranks; points represent average feature ranks; horizontal whiskers indicate the standard deviation across 20 resamples of the data.* ***(B)*** *Classifier performance (cross-validated training and validation performance) as a function of the number of features used; points represent average accuracies; vertical whiskers represent the standard deviation across 20 resamples of the data.*

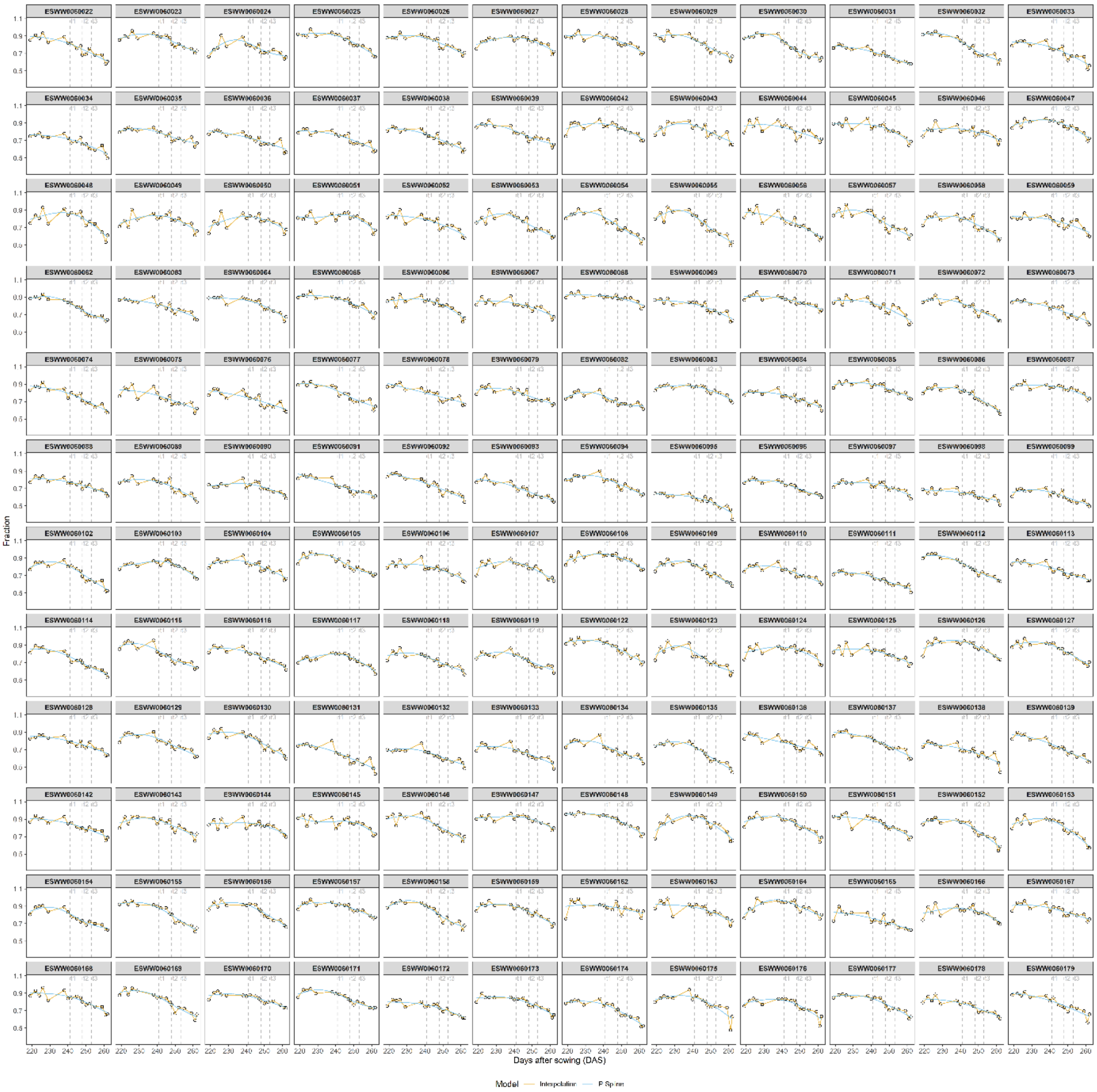


***Supplementary Figure S 12*** *Total vegetation cover; plot-level raw data extracted from image time series. The cover values on the y-axis are plotted against the time point of measurement, indicated in days after sowing, on the x-axis. Each facet represents raw data for one experimental plot (black circles, 17 measurement time points between heading and maturity). Plot time series were fitted using P-Splines. Please zoom in for details.*

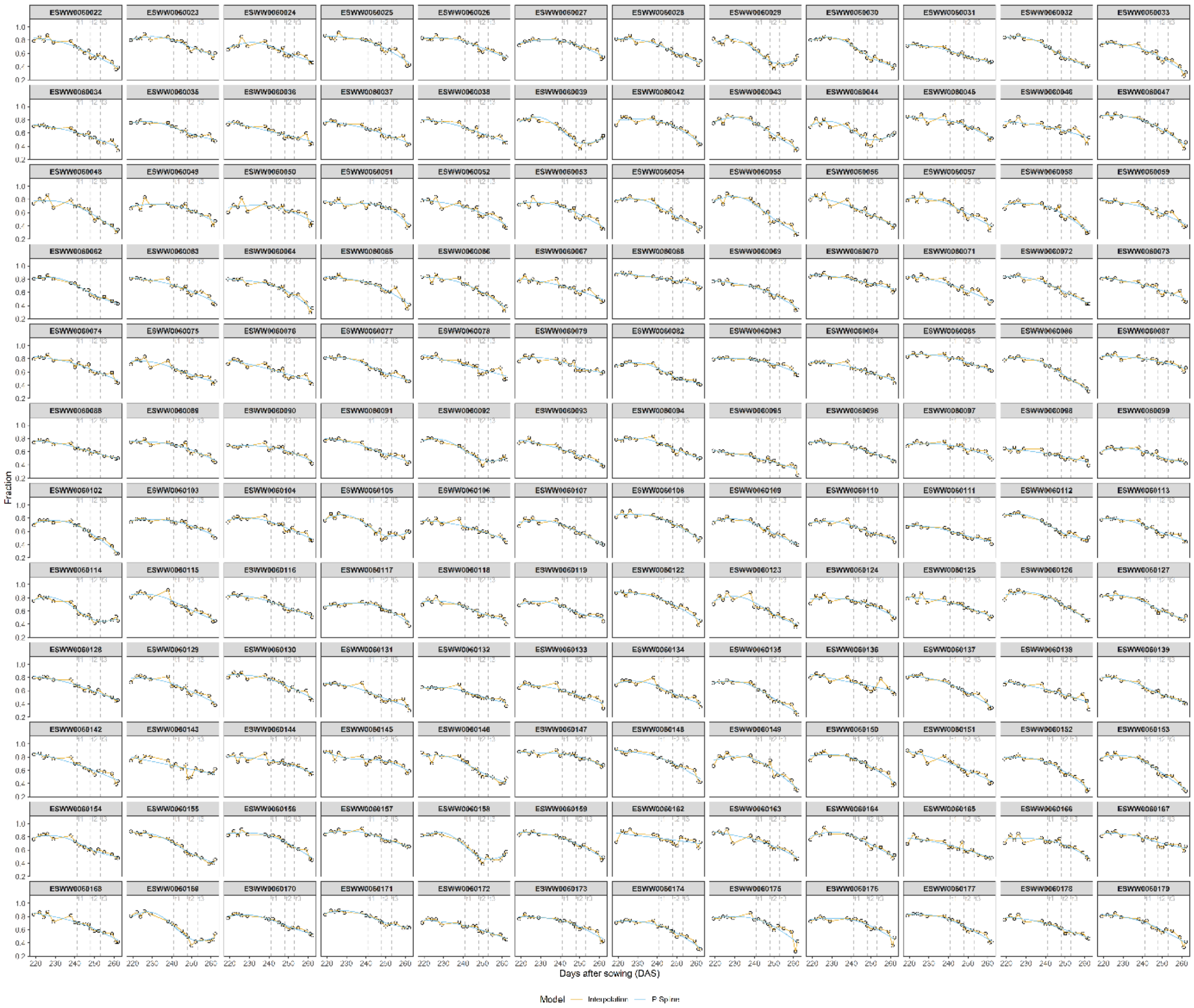


***Supplementary Figure S 13*** *Shoot (stem + leaves) cover; plot-level raw data extracted from image time series. The cover values on the y-axis are plotted against the time point of measurement, indicated in days after sowing, on the x-axis. Each facet represents raw data for one experimental plot (black dots, 17 measurement time points between heading and maturity). Plot time series were fitted using P-Splines. Please zoom in for details.*

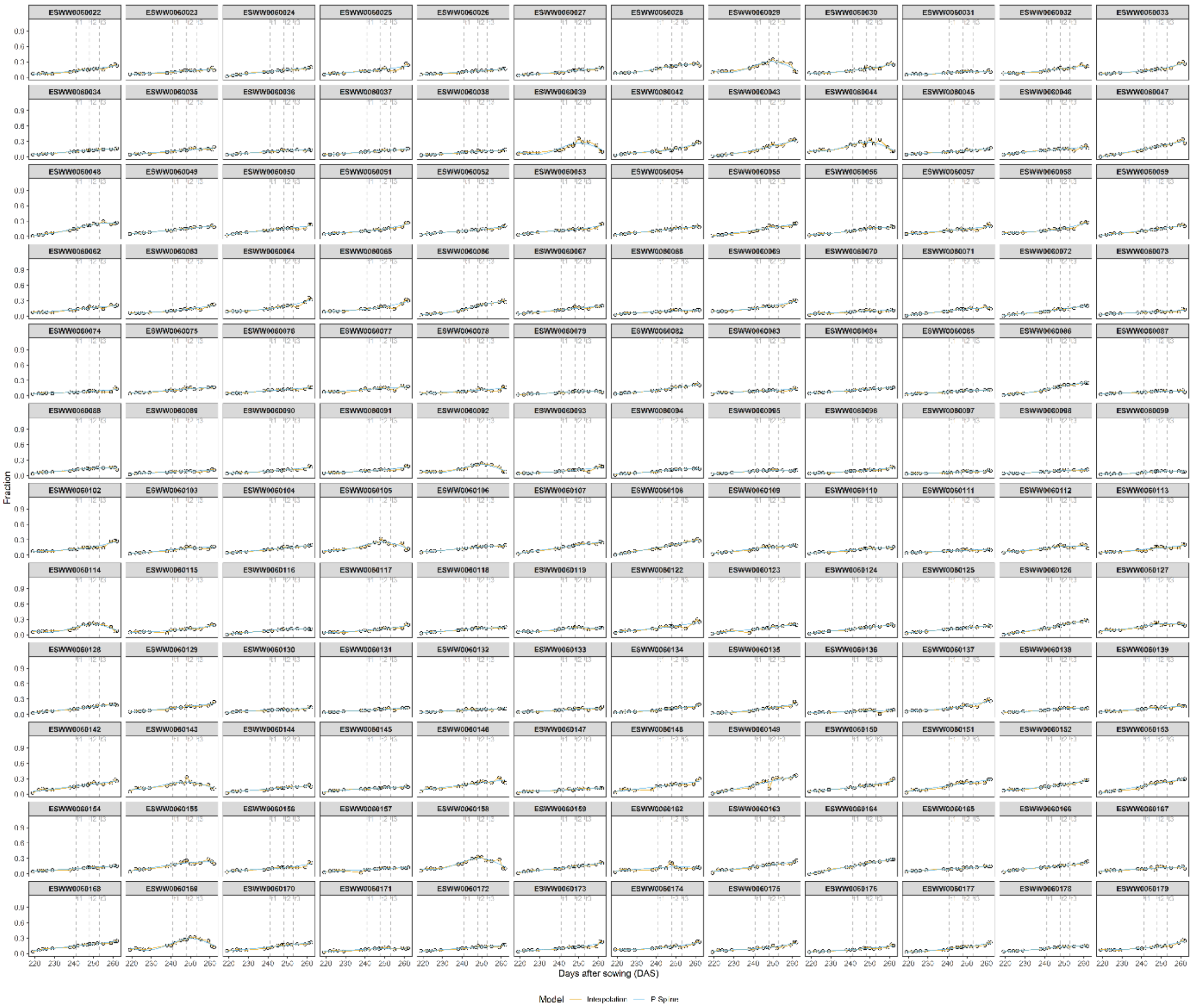


***Supplementary Figure S 14*** *Ear cover; plot-level raw data extracted from image time series. The cover values on the y-axis are plotted against the time point of measurement, indicated in days after sowing, on the x-axis. Each facet represents raw data for one experimental plot (black dots, 17 measurement time points between heading and maturity). Plot time series were fitted using P-Splines. Please zoom in for details.*

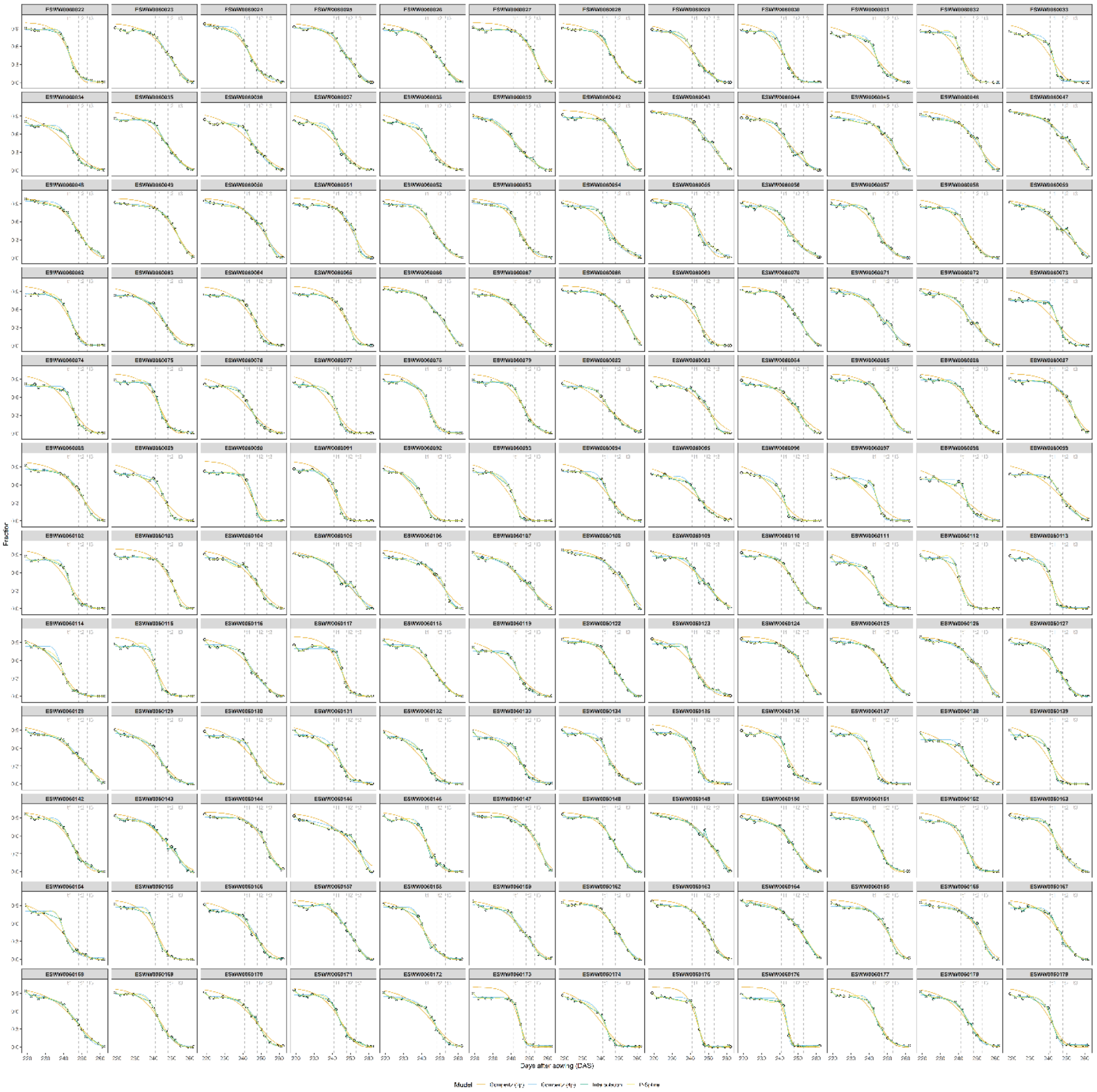


***Supplementary Figure S 15*** *Fraction of green healthy material in shoots (leaves + stems); plot-level raw data extracted from image time series. The cover values on the y-axis are plotted against the time point of measurement, indicated in days after sowing, on the x-axis. Each facet represents raw data for one experimental plot (black dots, 17 measurement time points between heading and maturity). Plot time series were fitted using different models, of which the four-parameter Gompertz model was finally selected based on visual inspection. Please zoom in for details.*

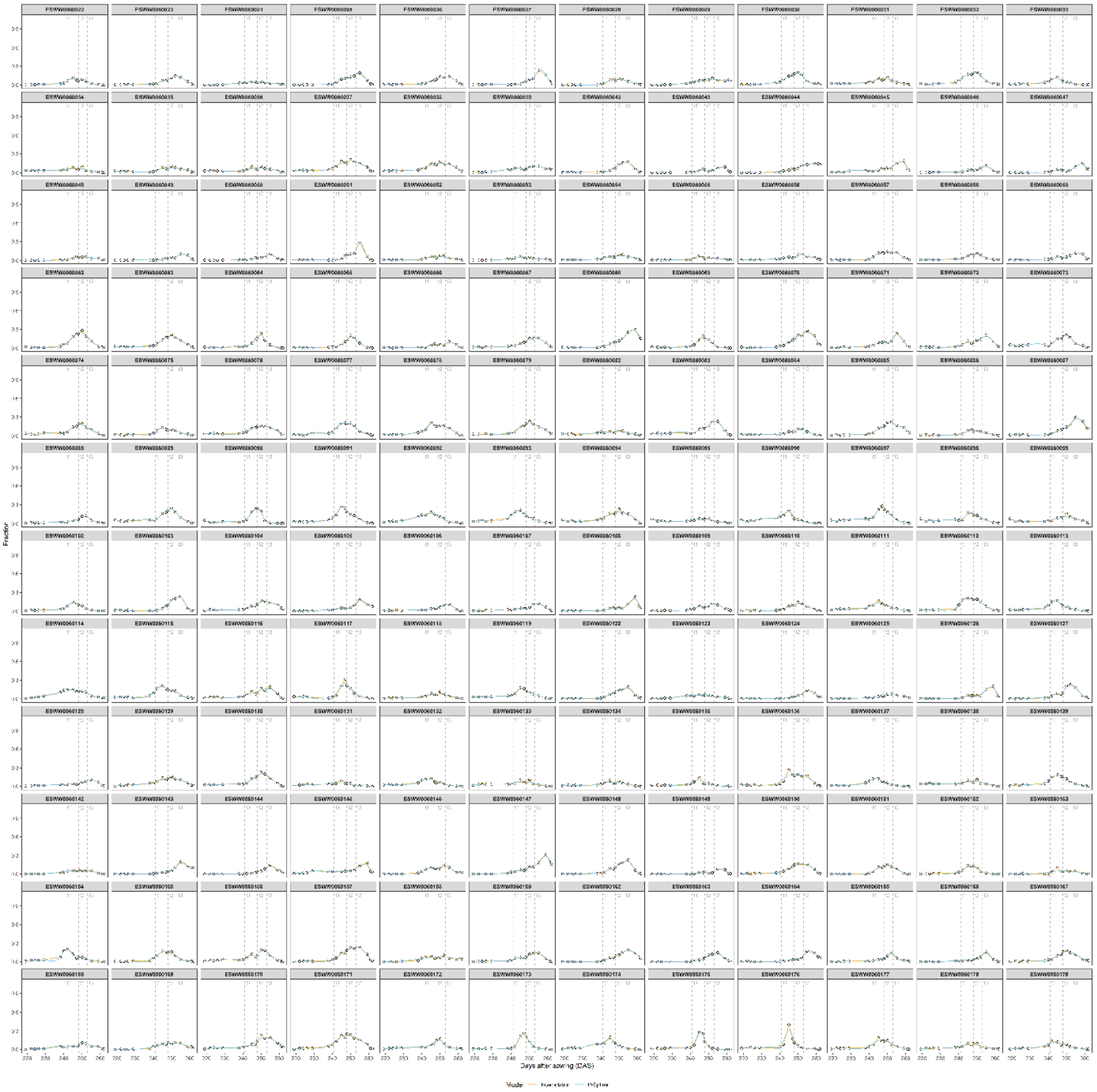


***Supplementary Figure S 16*** *Fraction of chlorotic material in shoots (leaves + stems); plot-level raw data extracted from image time series. The cover values on the y-axis are plotted against the time point of measurement, indicated in days after sowing, on the x-axis. Each facet represents raw data for one experimental plot (black dots, 17 measurement time points between heading and maturity). Plot time series were fitted using different P-Splines. Please zoom in for details.*

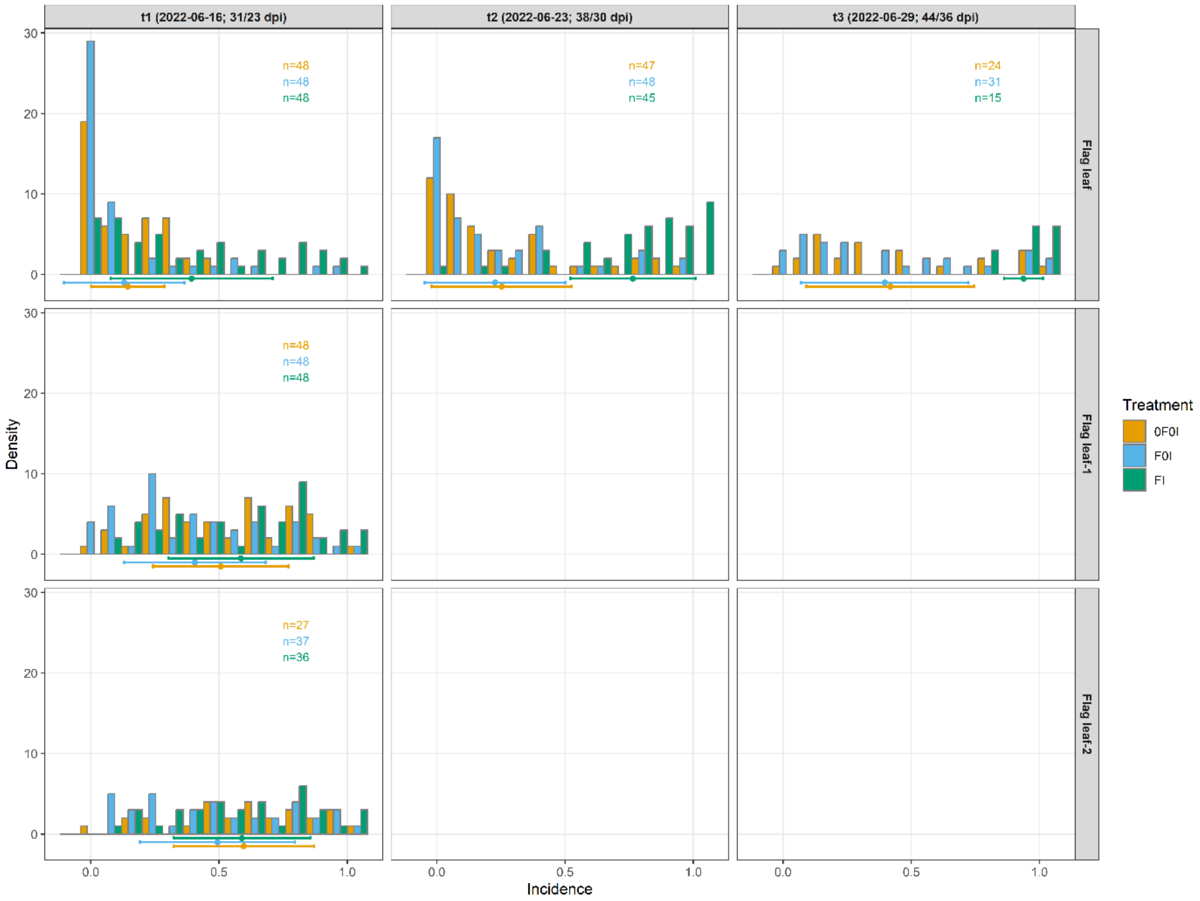


***Supplementary Figure S 17*** *STB incidence assessed by visually inspecting the flag leaves of 30 culms per plot. Assessments were made on the top-most three leaf layers at t1, and on flag leaves at t2 and t3. Points and horizontal whiskers below the bar plot represent per-treatment mean values and their standard deviations. Plot-based raw values are shown.*

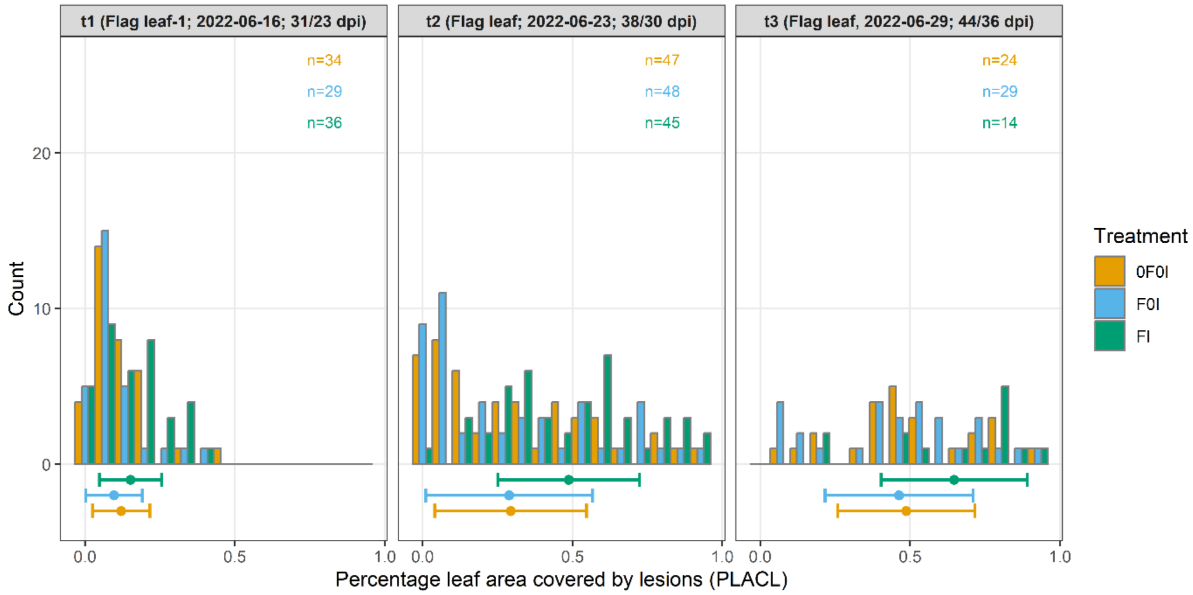


***Supplementary Figure S 18*** *Conditional STB severity assessed by estimating the percent leaf area covered by necrotic lesions for 8 infected leaves per plot. Assessments were made on the flag leaf minus one at t1, and on flag leaves at t2 and t3. Points and horizontal whiskers below the bar plot represent per-treatment mean values and their standard deviations. Plot-based raw values are shown.*

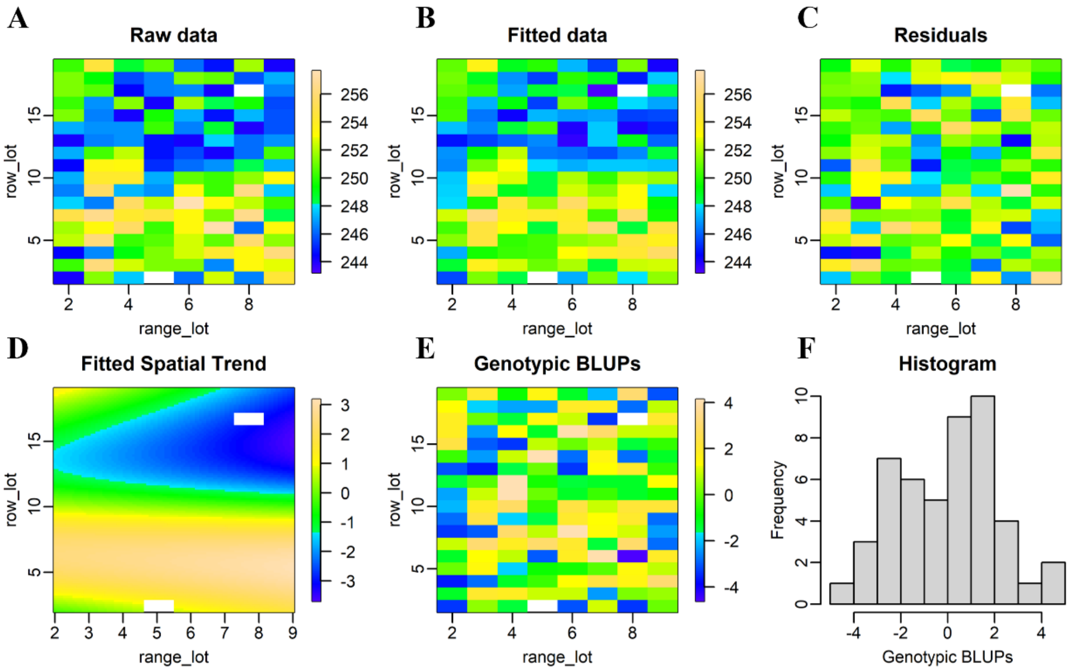


***Supplementary Figure S 19***. *Spatial patterns in the midpoint of senescence. The x-axis and y-axis represent the ranges and rows of the experiment, respectively.* ***(A)*** *Raw plot observations of the midpoint of senescence resulting from a linear interpolation of sequential visual scorings, indicated in days after sowing (DAS),* ***(B)*** *trait values fitted by the model,* ***(C)*** *residuals (observed – fitted),* ***(D)*** *fitted spatial trend, indicated in DAS,* ***(E)*** *genotypic estimates (best linear unbiased predictions BLUPs),* ***(F)*** *Histogram of the genotypic best linear unbiased estimators (BLUPS); here these values represent BLUPs for each genotype within each treatment.*

***Supplementary Table S 1*** *List of the wheat cultivars included in the field experiment. Cultivars were specifically selected to have similar phenology and final height but strongly contrasting canopy architectural and morphological traits, based on data from Anderegg et al. (2021). Specifically, the set comprised an equal number of cultivars with erect and planophile flag leaves and with high and low levels of flag leaf glaucousness.*

| GenName | FH | GS55 | Onsen | Awns | Fl0Ang | Fl0Glc | PLACL | TraitComb | Criterion | Originator | Accession Number | Recommended for |
|---|---|---|---|---|---|---|---|---|---|---|---|---|
| GULLIVER | 0.76 | 221 | 641 | 0 | 1 | 0 | 20.3 | LL | TraitComb | Nickerson International Research GEIE | na | GBR |
| KOBRA PLUS | 0.84 | 219 | 593 | 0 | 1 | 3 | 48.9 | LL | TraitComb | Hodowla Roslin Rolniczych Nasiona Kobierzyc Sp.z o. o. | RICP-0C0107039 | POL |
| HISTORY | 0.87 | 222 | 573 | 0 | 2 | 2 | 19.7 | LL | TraitComb | Bayerische Pflanzenzuchtgesellschaft eG & KG | RICP-01C0106532 | DEU |
| SCIROCCO | 0.92 | 218 | 587 | 0 | 5 | 2 | 32.1 | HL | TraitComb | Saatzuchtwirtschaft F. von Lochow-Petkus GmbH | RICP-01C0205033 | DEU |
| XENOS | 0.91 | 218 | 694 | 0 | 4 | 1 | 39.9 | HL | TraitComb | Fr. Strube Saatzucht | RICP-01C0106505 | AUT, LUX, DEU |
| TAIFUN | 0.85 | 216 | 656 | 0 | 5 | 3 | 37.8 | HL | TraitComb | Lochov-Petkus GMBH | K-57182; AUS-22191 | DEU, HUN, LTU, DNK |
| RAISON | 0.75 | 222 | 579 | 0 | 1 | 8 | 29.6 | LH | TraitComb | na | na | FRA |
| BRANDO | 0.71 | 222 | 586 | 0 | 1 | 8 | 41.8 | LH | TraitComb | Cambridge PB Twyford | K-64522 | GBR |
| POTENZIAL | 0.82 | 221 | 699 | 0 | 2 | 9 | 22.1 | LH | TraitComb | Deutsche Saatveredelung AG, OT Leutewitz | RICP-01C0107156 | DEU, CZE, DNK |
| DIAVEL | 0.86 | 218 | 600 | 0 | 5 | 8 | ? | HH | TraitComb | Agroscope/DSP (Federal Research Station for Agronomy) | na | CHE |
| RETRO | 0.82 | 222 | 545 | 0 | 4 | 8 | 19.6 | HH | TraitComb | Nickerson Limagrain GmbH | RICP-01C0106942 | DEU |
| BORNEO | 0.74 | 221 | 523 | 0 | 6 | 7 | 42.8 | HH | TraitComb | Saatzucht J.Breun GdbR | RICP-01C0106113 | DEU |
| FORNO | 0.78 | 220 | 584 | 0 | 3 | 4 | 40.4 | na | *Lr34* | Agroscope/DSP (Federal Research Station for Agronomy) | K-62003; AUS-23641 | CHE |
| ARINA | 0.77 | 220 | 494 | 0 | 1 | 7 | 22.4 | na | Resistant | Agroscope/DSP (Federal Research Station for Agronomy) | K-57737; K-57528; E-1014; AUS-21732 | CHE |
| MONTALBANO | na | na | na | 1 | na | na | na | na | Check | Agroscope/DSP (Federal Research Station for Agronomy) | na | CHE |
| AUBUSSON | 0.66 | 217 | 600 | 0 | 5 | 6 | 68.7 | na | Susceptible | Limagrain Verneuil Hold. | RICP-01C0107106 | FRA, ITA |

**GenName**: Cultivar name, **FH**: Final height, **GS55**: Heading date, indicated in days after sowing; **Onsen**: Onset of senescence, indicated in growing degree days after heading; **Awns**: Presence or absence of awns on ears, with '0' indicating absence and '1' indicating presence of awns, **Fl0Ang**: Visual scoring of flag leaf angle, with low values indicating erect flag leaves and high numbers indicating drooping flag leaves, **Fl0Glc**: Flag leaf glaucousness, with low values indicating low levels of glaucousness and high values indicating high levels of glaucousness, **PLACL**: Percent leaf area covered by lesions, data from Karisto et al. (2018), **TraitComb**: Trait combination represented by the cultivar, with 'L' and 'H' representing low and high values for Fl0Ang and Fl0Glc, respectively, **Criterion**: Selection criterion applied; either the trait combination, the presence of the *Lr34* gene, or the level of resistance. Information on the originator, accession number and cultivar recommendation were taken from the Genetic Resources Information System for Wheat and Triticale (GRIS), at URL http://wheatpedigree.net/, accessed on 28-02-2023. Trait data was based on a field experiment carried out in the wheat growing season of 2018/2019 at the Eschikon site (Anderegg et al., 2021). Assessments were made based on recommendations by Pask et al. (2012).

***Supplementary Table S 2*** *All pairwise correlations between the difference in STB severity and in vegetation fraction dynamics at the genotype-level (n = 16) as observed between the treatments 'FI' (early fungicide application + later inoculation – "clean STB") and 'F0I' (early fungicide application, no inoculation – "healthy control"). Trait contrasts between treatments at the genotype-level were calculated as the difference of the mean corrected plot values for each genotype in each treatment.*

| Level | Fraction | Parameter | Pearson r | p-value |
|---|---|---|---|---|
| shoot | chlr | max | -0.57* | 0.021074 |
| shoot | chlr | q1 | -0.566* | 0.022412 |
| veg | green | t20 | -0.552* | 0.026668 |
| shoot | chlr | h2 | 0.542* | 0.030131 |
| veg | chlr | q2 | 0.54* | 0.030941 |
| shoot | chlr | intq | 0.521* | 0.038416 |
| shoot | chlr | inth | 0.504* | 0.046292 |
| shoot | green | t20 | -0.478 | 0.060915 |
| shoot | chlr | q2 | 0.466 | 0.068723 |
| ear | chlr | Integral | 0.43 | 0.096718 |
| veg | chlr | max | -0.39 | 0.135335 |
| veg | green | M | -0.369 | 0.159356 |
| shoot | green | Integral | -0.366 | 0.163045 |
| veg | chlr | intq | 0.358 | 0.173551 |
| veg | chlr | h2 | 0.355 | 0.177089 |
| shoot | green | C | 0.355 | 0.177243 |
| shoot | chlr | h1 | -0.35 | 0.183292 |
| veg | green | Integral | -0.333 | 0.206873 |
| veg | green | t80 | -0.333 | 0.20756 |
| veg | chlr | q1 | -0.322 | 0.223976 |
| veg | green | A | 0.297 | 0.263584 |
| veg | green | t50 | -0.282 | 0.28923 |
| veg | chlr | inth | 0.273 | 0.307028 |
| ear | green | t80 | -0.267 | 0.317748 |
| shoot | green | M | -0.262 | 0.326241 |
| ear | green | Integral | -0.255 | 0.340279 |
| veg | green | b | 0.247 | 0.355477 |
| veg | chlr | h1 | -0.247 | 0.357007 |
| ear | chlr | max | 0.244 | 0.362922 |
| veg | green | C | -0.237 | 0.376781 |
| ear | green | t50 | -0.234 | 0.382958 |
| shoot | green | A | -0.184 | 0.494442 |
| ear | green | C | 0.183 | 0.498646 |
| ear | green | t20 | -0.172 | 0.523778 |
| ear | chlr | q2 | 0.167 | 0.53745 |
| shoot | green | t80 | -0.166 | 0.539583 |
| ear | green | A | -0.155 | 0.565653 |
| shoot | green | t50 | -0.154 | 0.569861 |
| shoot | chlr | Integral | -0.137 | 0.612169 |
| ear | chlr | h2 | -0.113 | 0.677868 |
| ear | chlr | h1 | -0.104 | 0.701744 |
| shoot | green | b | -0.099 | 0.713998 |
| ear | green | M | -0.086 | 0.750495 |
| ear | chlr | inth | -0.08 | 0.769334 |
| veg | chlr | Integral | -0.058 | 0.831249 |
| ear | chlr | q1 | -0.03 | 0.912789 |
| ear | chlr | intq | -0.024 | 0.929888 |
| ear | green | b | -0.003 | 0.991256 |
